# Maternal adaptations in mouse lactation are vulnerable to diet-induced excess adiposity

**DOI:** 10.1101/2023.06.30.547298

**Authors:** Jessica A. Breznik, Tatiane A. Ribeiro, Erica Yeo, Brianna K. E. Kennelly, Xuanyu Wang, Candice Quin, Braeden Cowbrough, Elena F. Verdú, Dawn M. E. Bowdish, Deborah M. Sloboda

**Affiliations:** Department of Medicine, McMaster Immunology Research Centre, McMaster University, Hamilton, Ontario, Canada; Michael G. DeGroote Institute for Infectious Disease Research, McMaster University, Hamilton, Ontario, Canada; McMaster Institute for Research on Aging, McMaster University, Hamilton, Ontario, Canada; Department of Biochemistry and Biomedical Sciences, McMaster University, Hamilton, Ontario, Canada; Farncombe Family Digestive Health Research Institute, McMaster University, Hamilton, Ontario, Canada; Firestone Institute of Respiratory Health, St Joseph’s Healthcare, Hamilton, Ontario, Canada; Department of Pediatrics, McMaster University, Hamilton, Ontario, Canada

**Keywords:** lactation, postpartum, gestational weight gain, postpartum weight retention, excess adiposity, obesity, high calorie diet, intestinal permeability, immunophenotype, monocyte, macrophage

## Abstract

Lifetime maternal risk of obesity is increased by excess gestational weight gain, which alters maternal metabolic adaptations in pregnancy, but the effects of excess weight retention/adiposity in the early postpartum period have not been extensively studied. The pathophysiology of excess adiposity and its accompanying immunometabolic dysregulation are associated with impaired intestinal barrier function in both non-pregnant and pregnant contexts. Using a mouse model, we investigated effects of diet-induced excess adiposity on maternal physiological adaptations during lactation. We report that in lactation, excess adiposity altered maternal intestinal morphology, influenced local immune cell composition and phenotype, increased intestinal permeability, and altered whole-body glucose metabolism as well as peripheral inflammation and immune cell composition. Many of these effects persisted two months post-lactation in mice with excess adiposity. Our findings have important implications for the development of interventions for periconceptual/perinatal excess adiposity and emphasize that further studies are needed to better understand effects of excess adiposity on maternal postpartum health.

## Introduction

Worldwide, over 40% of women gain excess weight during pregnancy^1,2^, which elevates risks of adverse maternal and fetal complications^3^. Even more women have postpartum weight retention^4,5^, which increases long-term risk of obesity and associated cardiometabolic conditions^6–8^. As the postpartum period of one pregnancy could be the preconception period of the next, excess weight gain and retention in one pregnancy may exacerbate risks to maternal and child health across successive pregnancies^9,10^. A better understanding of postpartum maternal physiology, especially in the context of excess adiposity, is thus necessary to inform the development of interventions to improve perinatal and lifelong maternal and child health.

Pregnancy and lactation require extensive and temporally-regulated adaptations to maternal physiology, metabolism, and immunity^11^, which necessitate adjustments in energy and nutrient absorption and expenditure^12^. Adaptations within the maternal intestines are important contributors to these metabolic demands, and include adjustment of tissue length and villus-crypt homeostasis^13–15^, increased vascularization and nutrient transport^13,15,16^, and changes to immune regulation^17^. There is also time-dependent modulation of maternal intestinal barrier function and the intestinal microbiota over the course of pregnancy^14,18^. As commensal microbes and microbial components and metabolites influence bone marrow hematopoiesis, these intestinal adaptations likely contribute to the gradual across-gestation modulation of peripheral immune cell composition and function towards a more myeloid-centric immunophenotype^19^, as well as changes in whole-body metabolism^20^. In pregnancy, the trajectory of maternal adaptations is influenced by maternal, fetal, and environmental signals, including maternal periconceptual/perinatal excess adiposity^14^.

An early effect of excess adiposity in non-pregnant mice is loss of gut barrier function^20^. The intestinal barrier has physical, biochemical, and immune components, including a single layer of constantly replenished epithelial cells, an overlying mucous layer formed by mucins and other bioactive molecules secreted by goblet cells, and immune cells that provide protection as well as promote tolerance^21^. Changes to villus and crypt structure, goblet cell numbers, and local macrophage populations, are implicated in excess adiposity-induced loss of intestinal homeostasis^,22–24^. Increased intestinal permeability is thought to augment translocation of luminal contents into the underlying mucosa and into the periphery, resulting in systemic inflammation and elevated circulating proinflammatory Ly6C^high^ monocytes, as well as the accumulation of monocyte-derived and tissue-resident proinflammatory macrophages within adipose tissue, which contribute to gradual metabolic dysfunction^25–27^. Therefore, alterations in gut barrier function and monocytes/macrophages have key roles in the pathophysiology of obesity in non-pregnant mice. Whether the same is true during lactation is unclear, although we have shown that intestinal barrier function is similarly vulnerable to excess adiposity during pregnancy^14^.

Maternal excess adiposity during pregnancy in rodent models alters intestinal epithelial structure^14^, expression of intestinal genes associated with nutrient transport, inflammation, and epithelial barrier integrity^28–31^, the commensal microbiome^18,31,32^, and increases intestinal permeability^14^. These changes may underlie reports of elevated circulating proinflammatory cytokines^33–35^ in pregnancies with excess adiposity as well as increased immunomodulatory factors such as bacterial lipopolysaccharide (LPS) and muramyl dipeptide (MDP)^28,36^. Excess adiposity in pregnancy in animal models and humans has accordingly also been reported to alter the composition and function of monocytes in peripheral blood, as well as macrophages in placental and adipose tissues^14,33,37,38^, and to dysregulate glucose, insulin, and fatty acid metabolism^39,40^. Thus, excess adiposity impacts intestinal structure and barrier function, and influences whole-body pregnancy adaptations.

No studies to date have investigated whether maternal excess adiposity also influences maternal adaptations in the early postpartum period. Herein, we used a mouse model to examine the potential modifying effects of excess adiposity in the periconceptual/perinatal period on maternal intestinal structure, function, and immune cell composition and phenotype, as well as peripheral metabolism and immunity, during lactation, and their persistence post-lactation.

## Methods

### Ethics

All animal procedures in this study were performed in accordance with Institutional Animal Utilization Protocols (AUP# 20-07-27). And were approved by the McMaster University Animal Research Ethics Board, following the recommendations of the Canadian Council for Animal Care.

### Animals

Wildtype C57BL/6J female and male mice five weeks of age were purchased from The Jackson Laboratory (cat#00064) and were maintained at the McMaster University Central Animal Facility. All mice were housed under specific pathogen-free conditions with constant ambient temperature (22°C) on a 12-hour light-dark cycle. Following one week of acclimation after arrival, female mice were cohoused two per cage.

All mice were initially maintained on a standard control chow diet (17% fat, 29% protein, 54% carbohydrates, 3.4 kcal/g; Harlan Teklad 22/5 Rodent Diet, #8640) with food and water provided *ad libitum*. Mating was performed by cohousing two female mice with a wildtype C57BL/6J male mouse (fed control chow diet). Upon confirmation of pregnancy (by weight gain between date of mating and 10-14 days later), female mice were individually housed and allowed to deliver spontaneously. Once a litter was observed, this was designated as postnatal day one, or P1. Since it is not unusual for C57BL/6J females to cannibalize their first litters, we generated second pregnancies in order to have viable litters for lactation.

First pregnancy litters were euthanized at P3, and one week later, female mice were allocated to either continue on the standard control chow diet or were placed on a high calorie diet (45% fat, 20% protein, 35% carbohydrates, 4.73 kcal/g, Research Diets Inc., #D12451), with equal body weight distribution between diet groups. Following two weeks of dietary intervention, female mice were mated with a wildtype male mouse (fed control chow diet) at the end of the light cycle. The same male-female mouse pairing was used for all first and second pregnancy mating. Successful mating was identified the next morning by the presence of a vaginal plug (embryonic day E0.5 of gestation). Upon confirmation of pregnancy, female mice were allowed to deliver spontaneously. Pregnant mice were maintained on the control chow or high calorie diet throughout pregnancy, lactation, and post-weaning/lactation. Mice were checked daily after E18.5 for the presence of a litter.

Following second pregnancies, litters were standardized to six pups per dam at P7, with an equal sex ratio when possible, during lactation. Dams with litters of less than five pups were excluded from the study. As summarized in Figure 1, assays were performed on four independent cohorts during lactation between P21-P23, within 24 hours of weaning. One cohort was used to examine outcomes ∼1.5-2 months post-lactation, including assessments of oral glucose tolerance (∼P60), peripheral blood immunophenotyping (∼P67), intestinal permeability (∼P73), as well as circulating cytokines and intestinal histomorphology (∼P80).

**Figure 1.**
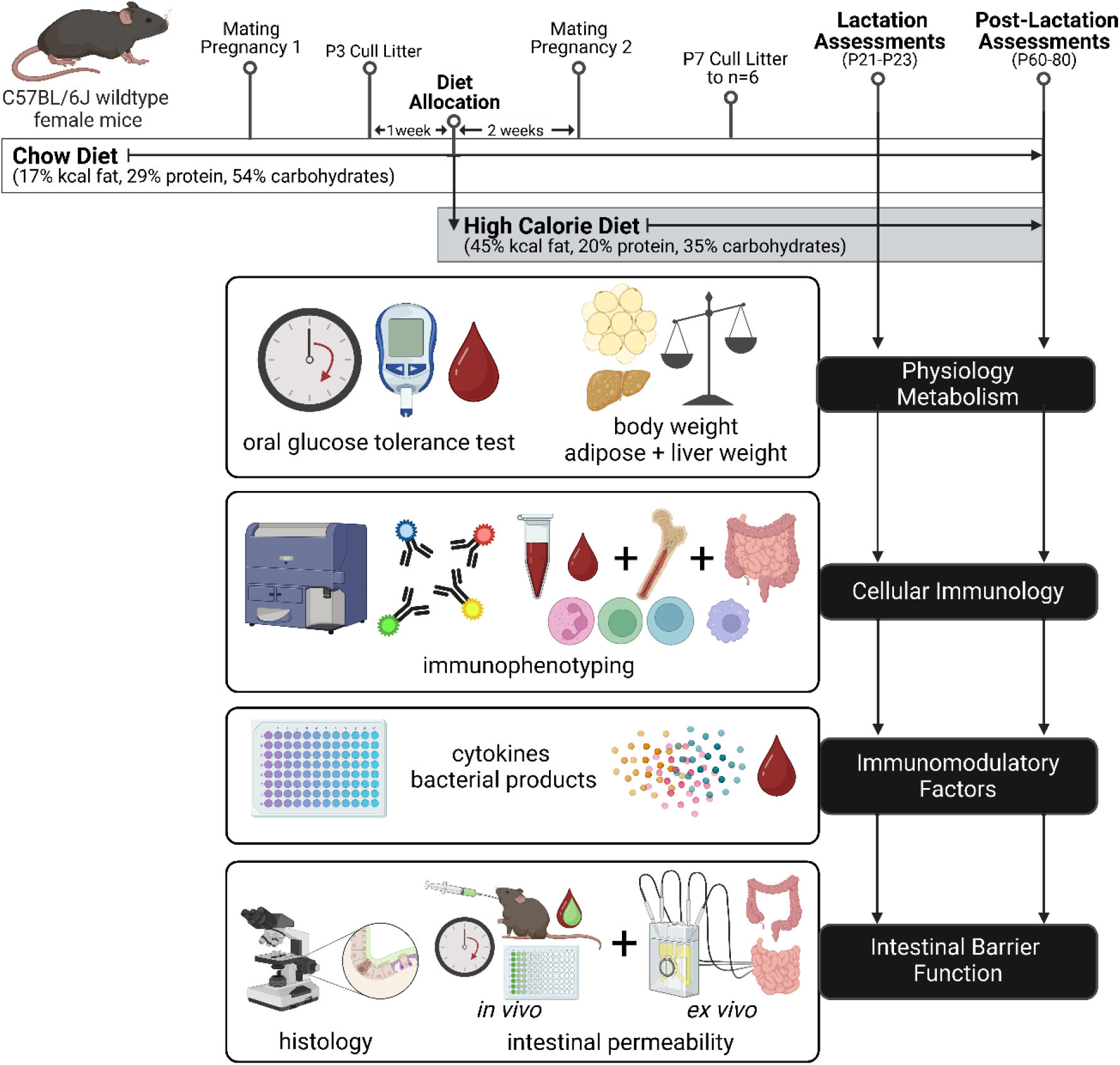
Study overview.

### Body weight, caloric intake, fasting glucose, and fasting insulin

Mouse body weight and/or food intake were monitored weekly pre-pregnancy and at E0.5, E6.5, E10.5, E14.5, and E18.5, as well as at P7, P14, P21 in lactation, and at P28, P35, P42, P49, and P56 post-lactation. Caloric intake was calculated based on the amount of food consumed and individual mouse weight per experiment cage ([(food eaten/number of mice in cage)*(g/kcal food energy density)]/mouse body weight). Whole-body EchoMRI imaging (Bruker Minispec LF90-II) was used to determine percent body fat of total mouse weight.

Blood glucose concentrations were measured with the MediSure Blood Glucose Meter system glucometer and test strips (Bowers Medical Supply) after a 6 hour fast (3 am – 9 am), or in the morning without fasting, via tail vein nick. To collect serum for analysis of insulin levels, blood was collected with a heparinized capillary tube via tail vein after a 6 hour fast (3 am – 9 am), centrifuged at 8000 rpm for 10 minutes, and then the supernatant was removed and frozen at −80°C until analysis. Insulin concentrations were measured using a commercial ELISA kit following the manufacturer’s instructions (AIS Toronto Biosciences, #32380). Assay results were recorded using the BioPlex 200 system and BioPlex Manager software v6.1 (Bio-Rad Laboratories, Inc.). Gonadal fat, mesenteric fat, and liver weights were recorded at endpoint.

### Oral Glucose Tolerance Test

Glucose (Sigma-Aldrich, #G5767; 2g/kg) was orally administered to mice by gavage after a 6 hour fast (3 am – 9 am). Blood glucose was measured via tail vein prior to glucose administration (0 min) and after 15, 20, 30, 40, 60, 90, and 120 min, using the MediSure Blood Glucose Meter system glucometer and test strips (Bowers Medical Supply).

### Assessments of circulating bacterial products

Peripheral blood was collected retro-orbitally with heparinized capillary tubes under isoflurane anesthesia, centrifuged at 8000 rpm for 10 minutes, and serum was aliquoted and frozen at −80°C until analysis. Muramyl dipeptide and lipopolysaccharide were quantified in serum using in-house colourimetric reporter bioassays^41^. HEK-Blue TLR4 cells (Invivogen, #hkb-htlr4) and HEK-Blue-mNOD2 cells (Invivogen, #hb-mnod2) were transfected with the pNifty-SEAP reporter plasmid (Invivogen, #pnifty2-seap), containing NFκB transcription factor binding sites and a secreted alkaline phosphatase (SEAP) reporter gene, and cultured 37°C in a 5% CO_2_ tissue culture incubator in complete DMEM (DMEM with 10% (v/v) FBS, 1% (v/v) penicillin-streptomycin, and 1% (v/v) L-glutamine), supplemented with blasticidin and zeocin. Cells were seeded in a 96-well place at a density of 4×10^3^ cells per well and incubated overnight. Serum samples were diluted 1:5 in PBS, then 1:1 in sterile water, and heat inactivated at 75°C for 5 minutes. Media was removed prior to the addition of heat-inactivated serum diluted 1:20 in HEK Blue Detection media (Invivogen, #hb-det3). Cells were incubated for 48 hours and then absorbance was measured at 630 nm with a plate reader (SpectraMax). Standard curves of MDP (Invivogen, #tlrl-mdp) and LPS (Invivogen, #tlrl-pb5lps) were included to calculate relative amounts of MDP and LPS from absorbance units, after subtraction of background fluorescence of wells with cells and media in the absence of serum. Samples were prepared in triplicate and the assay was performed in duplicate.

### Serum cytokine measurements

Serum cytokines were measured using a U-PLEX Custom Biomarker Group 1 (mouse) Assay Meso Scale Discovery kit from Meso Scale Diagnostics (K15069L-1), on a 1300 Meso® QuikPlex SQ 120MM, following the manufacturer’s guidelines. Assessed cytokines (lower limit - upper limit of detection in pg/mL): TNF (1.3 – 6200), IFN-γ (0.16 – 2900), IL-1β (3.1 – 13000), IL-6 (4.8 – 16000), IL-10 (3.8 – 22800), MCP-1 (1.4 – 1400).

### *In vivo* and *ex vivo* assessments of intestinal permeability

*In vivo* assessment of whole-intestine permeability was performed as previously described^22^. Mice were fasted for 4 hours in the morning (6 am – 10 am) and during the assay. In brief, tail blood was collected at baseline and 30, 60, 90, 120 and 240 minutes after oral gavage with 50 mg/kg body weight 4 kDa fluorescein-isothiocyanate-conjugated dextran (FITC-dextran; Sigma-Aldrich, #46944) diluted to 8mg/mL concentration in PBS (pH 7.4; 1.8 mM KH_2_PO_4_, 2.7 mM KCl, 10 mM Na_2_HPO_4_, 137 mM NaCl).

*Ex vivo* assessments of gut barrier function were performed with sections of distal ileum and proximal colon using Ussing chambers as previously described^22^. Short-circuit current response (Isc) was used to measure net active ion transport after ∼30 minutes equilibration. Tissue conductance (G; mS/cm^2^) was calculated from measurements of potential difference and short circuit current by Ohm’s law. Paracellular permeability was assessed by mucosal-to-serosal flux (%flux/cm^2^/hr) of the small inert probe ^51^Cr-EDTA (360 Da; 6 μCi/mL) (Perkin Elmer), by taking an initial radiolabelled sample of mucosal buffer and then samples from the serosal compartment every 30 minutes for 2 hours, with assessment in a liquid scintillation counter (LS6500 Multi Purpose Scintillation Counter, Beckman Coulter, Inc.).

### Quantification of leukocytes from blood, bone marrow, and intestinal tissues

Peripheral blood was collected retro-orbitally with heparinized capillary tubes under isoflurane anesthesia in the morning to minimize effects of diurnal variation^42^. Whole blood was stained as previously described for quantitation of monocytes and neutrophils, NK cells, B cells, CD4^+^ T cells, and CD8^+^ T cells^43^. Bone marrow was extracted from femurs and prepared as previously described for assessment of monocytes^27^. Intestinal ileum and colon tissues were removed and processed for flow cytometry as published^22^. All fluorophore-conjugated antibodies used in immunophenotyping are specified in Supplementary Table 1. Data were analyzed using FlowJo software (v10.8; Tree Star). Whole blood and intestinal tissue leukocytes were identified as previously published^22,43^, and bone marrow leukocytes were analyzed as shown in Supplementary Figure 1. CountBright Absolute Counting Beads (Life Technologies, #C3650) were used to determine absolute cell numbers. Surface expression was quantified for each fluorescence marker by measuring geometric mean fluorescence intensity and subtracting background geometric mean fluorescence intensity of isotype or fluorescence-minus-one controls.

### Intestinal tissue measurements and immunohistochemistry

The intestine was collected from stomach to rectum, mesenteric fat was removed, and the intestine was allowed to lay flat. Small intestine length was measured from the pylorus to the cecum. Colon length was measured from the cecum to the rectum. Caecal weight was determined. Assessment of gross morphological traits of villus and crypt structure, as well as goblet cell counts, were performed following established protocols^14^. In brief, tissue sections of 1 cm were collected from the distal ileum and distal colon, fixed in Carnoy’s fixative, embedded in paraffin, cut in 4 μm sections, stained with Periodic Acid-Schiff (PAS) (Sigma-Aldrich, #395B-1KT) to identify neutral mucins, and counterstained with Gill’s Hematoxylin No. 3 (Sigma-Aldrich, #GHS332-1L) to identify nuclei. Images were captured using a Nikon Eclipse NI microscope (Nikon Eclipse NI-S-E, #960122) and Nikon DS-Qi2 Colour Microscope Camera at 20x magnification, and analyzed with the Nikon NIS Elements Imaging Software (v4.30.02). Four tissue sections per dam, and 20 villi and 20 crypts per section, were assessed to determine average villus length and crypt depth. Four tissue sections per dam, and 10 villi and 10 crypts per tissue section, were assessed to determine average goblet cell counts per villus and crypt.

### Statistical analysis

The dam (i.e., female mouse, not offspring) was the biological replicate in all analyses. Data from age-matched virgin (nulligravid/never-pregnant) female mice used for comparisons was obtained from published datasets^14,27^. Data were analyzed and plotted with GraphPad Prism (v9.4.0; GraphPad Software) or RStudio (v2023.03.0+386) using packages and default functions from CRAN (https://cran.r-project.org/). Comparisons between chow-fed and high calorie-fed mice in lactation or post-lactation were performed by unpaired two-tailed Student’s t test (parametric) with Welch’s correction (for unequal variances or sample size) or Mann-Whitney U test (non-parametric) according to normality. Comparisons between never-pregnant female mice and mice in lactation and post-lactation were performed by one-way ANOVA (parametric) with the Tukey correction for multiple comparisons, or Kruskal-Wallis test (non-parametric) with Dunn’s correction for multiple comparisons. Comparisons of litter sizes and weights by biological sex and diet group were assessed by two-way ANOVA, with post-hoc multiple comparison tests with Tukey’s correction. Two-way repeated measures ANOVA with Šidák’s post-hoc test was used to assess oral glucose tolerance and intestinal permeability by FITC assay between diet groups. Permutational multivariate analysis of variance (PERMANOVA) with the *adonis2* function (*vegan* package) was used to examine differences in immune cell population composition. The permutation test for homogeneity of multivariate dispersions (PERMDISP), with the *betadisper* and *permutest* functions (*vegan* package), was used to compare the median distance-to-centroid of immune cell population variation within experimental groups. The *prcomp* function and *ggbiplot* package were used to generate Principal Component Analysis (PCA) ordination plots on PC1 and PC2, after mean centering and variance scaling. Statistical significance was defined as a p value of < 0.05.

### Data availability

All experimental data supporting the findings of this study are available from figshare: https://figshare.com/s/b2fb4902df727238a1ab.

## Results

### High calorie diet induces maternal excess adiposity and metabolic dysregulation during lactation

Female mice were fed a standard chow diet (CTL; 17% kcal fat) or high calorie diet (HC; 45% kcal fat) for two weeks prior to and throughout second pregnancies, lactation, and post-lactation to endpoint (Figure 1). Maternal body weight, cumulative weight gain, and energy consumption were monitored throughout the study (Supplementary Figure 2). Litter sizes were adjusted to a maximum of six offspring at P7, with equal sex ratios when possible, as litter size influences maternal intestinal physiological changes during lactation^13^. Prior to adjustment of litter size at P7, total litter sizes and sex ratios were similar between diet groups (Supplementary Figure 3a-b; Supplementary Table 2). Offspring weights were similar between diet groups and by sex at P7, but there was a main effect of maternal diet on offspring weights at P14 and P21 (Supplementary Figure 3c).

At the lactation endpoint (P21-23), mice fed the HC diet had ∼30% greater weight gain from the start of diet allocation compared to mice fed the CTL diet (mean ± SD at P21, HC mice: 6.51 ± 2.95 g; CTL mice: 4.59 ± 2.38 g) (Figure 2a; Supplementary Figure 2b). This increase in body weight was apparent despite total energy consumption (assessed as kilocalories per gram body weight) not being consistently higher in HC-fed mice compared to CTL-fed mice, from the start of diet allocation through to P21 (Supplementary Figure 2c). HC-fed mice also had excess adiposity, indicated by increased fat relative to whole-body weight (Figure 2b) and higher gonadal fat mass (Figure 2c), though liver weight was decreased (Figure 2d). Diet-induced excess adiposity in the HC-fed mice was also accompanied by increased fasting insulin (Figure 2e) and elevated fasting and non-fasting blood glucose levels (Figure 2f). There was a main effect of diet on oral glucose tolerance, but this difference did not reach statistical significance by area under the curve (Figure 2g). Therefore, a periconceptual/perinatal high calorie diet had no effects on offspring number or sex ratio but induced excess maternal weight gain and adiposity, and maternal hyperglycemia and hyperinsulinemia, during lactation.

**Figure 2.**
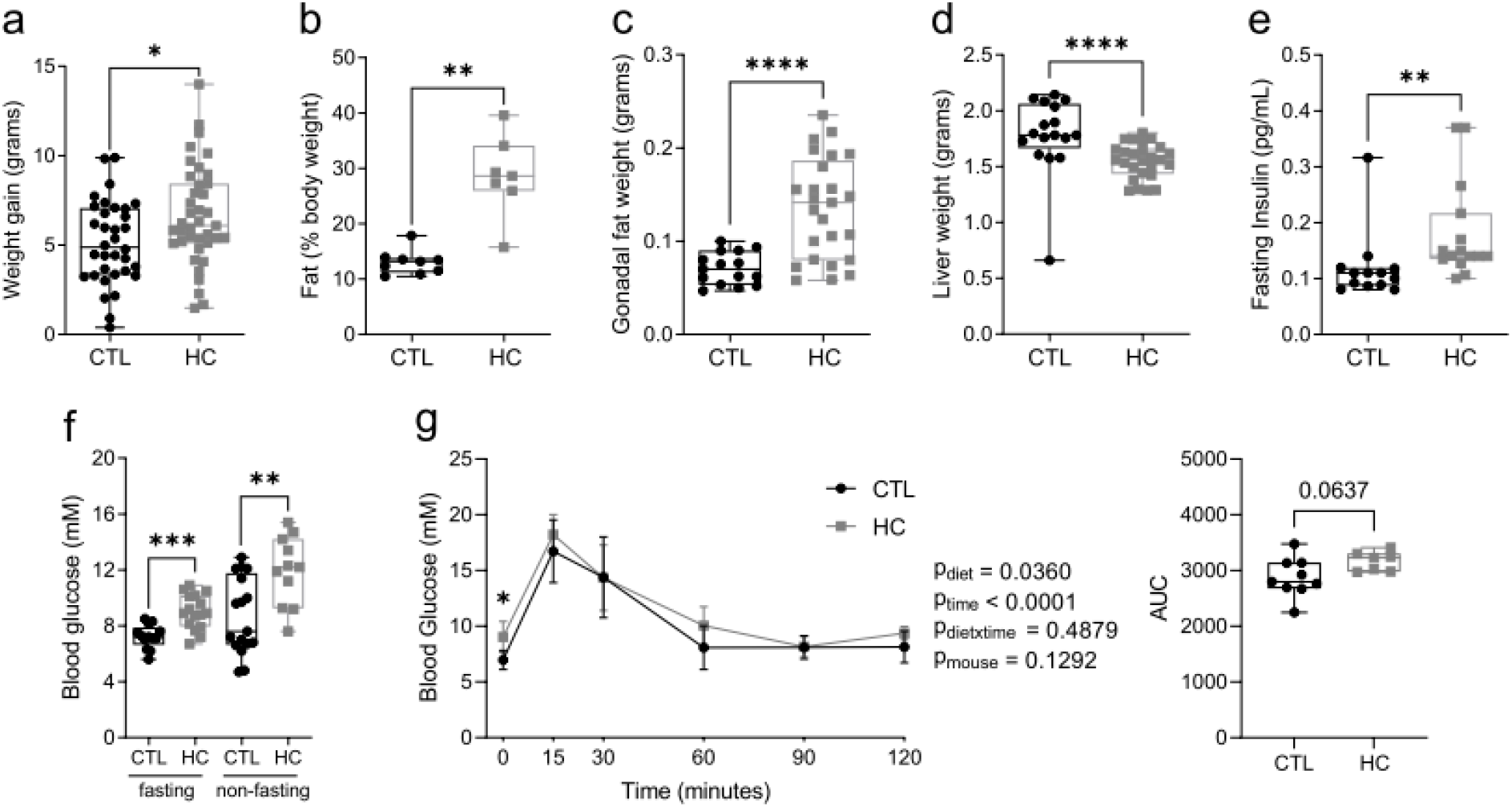
High calorie diet induces excess adiposity and alters maternal metabolism in lactation. Maternal physiological parameters were measured during lactation in mice fed chow (CTL) or high calorie (HC) diet. (**a**) maternal weight gain from start of diet allocation. (**b**) fat as a proportion of total body weight calculated from EchoMRI imaging. (**c**) gonadal fat weight. (**d**) liver weight. (**e**) fasting blood insulin. (**f**) blood glucose – fasting and non-fasting. (**g**) oral glucose tolerance test and area under the curve (AUC). Data in **a**-**f** and AUC in **g** are shown as box and whisker plots, minimum to maximum, with the center line at the median. Data in **g** are shown with a dot at the mean with error bars at ± standard deviation. Each data point indicates one mouse, with CTL as black circles and HC as grey squares. For **a** and **c**-**f**, CTL n=13-34, HC n=11-39. For **b** and **g**, CTL n=9, HC n=7. Statistical significance was assessed in **a**-**f** and AUC in **g** by two-tailed Student’s t test with Welch’s correction for unequal variances or Mann-Whitney U test according to data normality, and in **g** by two-way repeated measures ANOVA with Šidák’s post-hoc test. *p<0.05, ***p<0.01,*** p<0.001, ****p<0.0001.

### High calorie diet and excess adiposity alter maternal intestinal physiology during lactation

Altered intestinal structure and histomorphology are known to be associated with diet-induced excess adiposity in non-pregnant mice and in pregnancy^23,24^. We found that a HC diet induced excess mesenteric adiposity during lactation (Figure 3a). The small intestines were shorter in HC-fed mice compared to CTL-fed mice (Figure 3b; mean ± SD, CTL: 44.8 ± 1.0 cm; HC: 40.8 ± 3.5 cm), as were colon lengths (Figure 3c: CTL: 9.0 ± 1.4 cm; HC: 7.3 ± 0.7 cm). Cecal weight was 2.7-fold lower in HC-fed mice compared to CTL-fed mice (Figure 3d). Ileum villus height (Figure 3e) and crypt depth (Figure 3f) were decreased in mice fed HC diet. The numbers of goblet cells per ileum villus (Figure 3g) and crypt (Figure 3h) were similar in CTL-fed and HC-fed mice. Colon crypt depth (Figure 3i) and goblet cell numbers per crypt (Figure 3j) were decreased in HC-fed mice compared to CTL-fed mice. Representative images of PAS and hematoxylin staining are shown in Figure 3k. In summary, a periconceptional/perinatal HC diet and excess adiposity during lactation had significant tissue-specific effects on intestinal structure, morphology, and goblet cell numbers.

**Figure 3.**
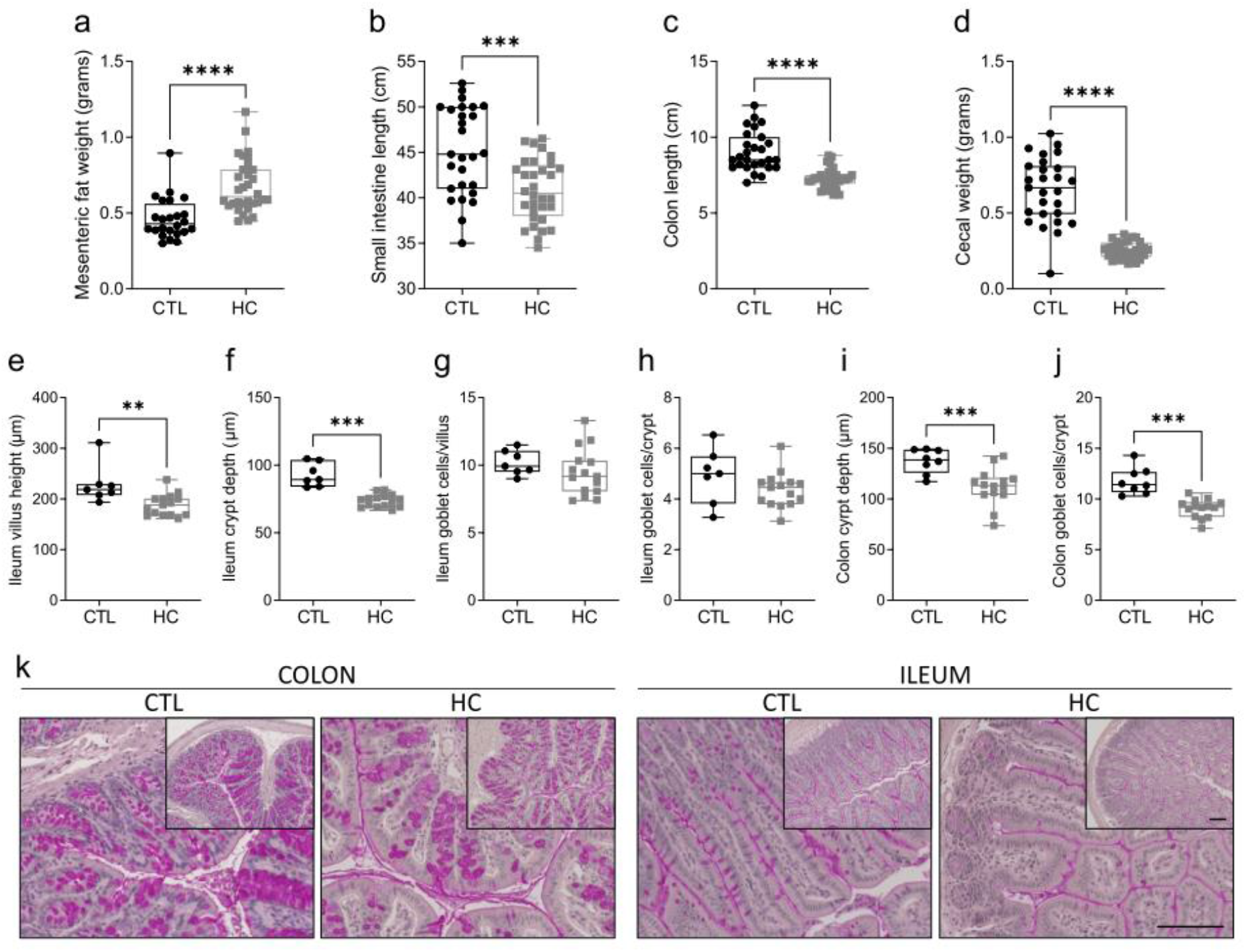
High calorie diet and excess adiposity alter maternal intestinal histomorphology in lactation. Maternal intestinal physiological parameters were measured during lactation in mice fed chow (CTL) or high calorie (HC) diet. (**a**) mesenteric fat weight. (**b**) small intestine length. (**c**) colon length. (**d**) cecal weight. (**e**) ileum villus height. (**f**) ileum crypt depth. (**g**) ileum goblet cells per villus. (**h**) ileum goblet cells per crypt. (**i**) colon crypt depth. (**j**) colon goblet cells per crypt. (**k**) representative panels of PAS and hematoxylin staining of ileum and colon sections taken at 20x magnification with 10x inset. Scale bars: 100 µm. Data in **a**-**j** are presented as box and whisker plots, minimum to maximum, with the center line at the median. Each data point indicates an individual mouse with CTL as black circles and HC as grey squares. For **a**-**d**, CTL n=24-27 and HC n=32-33. For **e**-**h**, CTL n=7 and HC n=15. For **i** and **j**, CTL n=8 and HC n=14. Statistical significance was assessed by two-tailed Student’s t test with Welch’s correction for unequal variances or Mann-Whitney U test according to data normality. **p<0.01, ***p<0.001, ****p<0.0001.

### High calorie diet and excess adiposity increase maternal intestinal permeability during lactation

The changes that we observed to intestinal structure in HC-fed mice may impact intestinal barrier function. An *in vivo* assay was initially used to assess whole-intestine permeability, by measuring plasma fluorescence levels after oral gavage of FITC-dextran. We observed that there was a main effect of diet, and an increase in whole-intestine permeability (measured by area under the curve), in HC-fed mice compared to CTL-fed mice during lactation (Figure 4a). To provide a more sensitive and location-specific assessment of intestinal barrier function, an *ex vivo* culture assay (using Ussing chambers) with tissues from the distal ileum and proximal colon was used to measure ion secretion and paracellular permeability (i.e., movement between adjacent cells). Measurements of ion short current (i.e., active transport of ions) were similar in ileum tissues of HC and CTL-fed diet groups but increased in colons of HC-fed mice compared to CTL-fed mice (Figure 4b). The increase in colon ion secretion may indicate an increase in chloride ion flux and altered water movement^44^. Conductance was higher in ileum tissues of HC-fed mice compared to CTL-fed mice, but comparisons between the dam diet groups were similar in the colon (Figure 4c). Conductance provides an indication of ion flux, dependent on changes in tight junction protein expression and function^44^. Mucosal-to-serosal movement of ^51^Cr-EDTA, an assessment of paracellular permeability (and an indication of tight junction function)^44^, was higher in both the ileum and colon tissues of HC-fed mice compared to CTL-fed mice (Figure 4d). Therefore, HC diet-induced excess adiposity impaired maternal intestinal barrier function during lactation.

**Figure 4.**
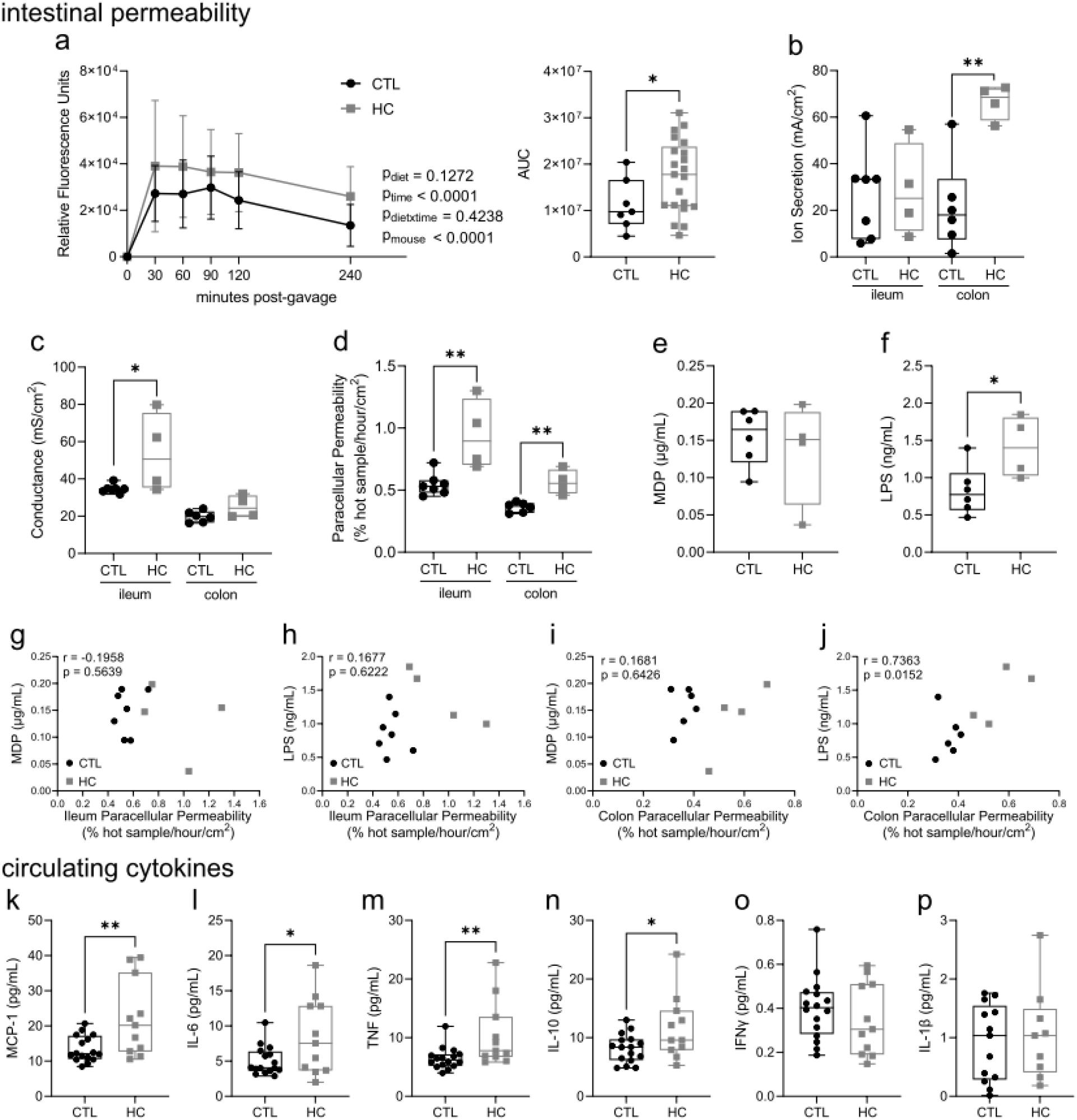
High calorie diet and excess adiposity increase maternal intestinal permeability and systemic inflammation in lactation. Maternal assessments were performed during lactation in mice fed chow (CTL) or high calorie (HC) diet. (**a**) *in vivo* whole-intestine permeability measured by FITC fluorescence in plasma post-gavage and area under the curve (AUC). *Ex vivo* assessments of proximal ileum and distal colon tissues: (**b**) ion secretion measured by ion short current, (**c**) tissue conductance, (**d**) paracellular permeability measured by mucosal-to-serosal flux of Cr^51^-EDTA. Circulating bacterial products (**e**) MDP and (**f**) LPS. Correlations of MDP and LPS with paracellular permeability in the ileum (**g**-**h**) and colon (**i**-**j**). Circulating cytokines: (**k**) MCP-1, (**l**) IL-6, (**m**) TNF, (**n**) IL-10, (**o**) IFNγ, (**p**) IL-1β. Data in **a** are shown with dots at the mean ± standard error of the mean. Data for AUC in **a** and data in **b**-**f** and **k-p** are shown as box and whisker plots, minimum to maximum, with the center line at the median. Each data point in AUC in **a** and **b-p** indicates an individual mouse, with CTL as black circles and HC as grey squares. For **a**, CTL n=7 and HC n=21. For **b**-**j**, CTL n=6-7 and HC n=4. For **k**-**p**, CTL n=16 and HC n=11. Statistical significance in **a** was assessed by two-way repeated measures ANOVA with Šidák’s post-hoc test, for AUC in **a**, **b**-**f**, and **k**-**p** by two-tailed Student’s t test with Welch’s correction for unequal variances or Mann-Whitney U test, and in **g**-**j** by Pearson’s correlation. *p<0.05, **p<0.01.

Increased paracellular permeability could lead to translocation of microbial components into systemic circulation, as reported in non-pregnant and late pregnancy mice with excess adiposity^28,36^, which may contribute to our observed changes in maternal metabolism (Figure 2e-f). We measured bacterial cell wall components MDP and LPS in serum (Figure 4e-f) and assessed their relationship with paracellular permeability (Figure 4g-j). There was an increase in circulating LPS in the HC-fed mice (Figure 4f), and a significant correlation between circulating LPS and colon paracellular permeability (Figure 4j). We also considered that this impaired intestinal barrier function may be accompanied by systemic inflammation. HC-fed mice with excess adiposity had elevated serum concentrations of MCP-1, IL-6, and TNF (Figure 4k-m), as well as IL-10 (Figure 4n), though IFNγ and IL-1β concentrations (Figure 4o-p) were similar between diet groups. Therefore, by combining measurements of intestinal morphology and serum immunomodulatory factors with *in vivo* and *ex vivo* measurements of intestinal permeability, our data provide clear evidence of elevated intestinal permeability in lactation in mice with HC diet-induced excess adiposity, which is accompanied by systemic inflammation.

### High calorie diet and excess adiposity alter maternal intestinal immune cell population dynamics during lactation

As changes to intestinal physiology and compromised barrier function influence local immunity and *vice versa*^20^, we next assessed tissue immune cell composition in the ileum and colon. In the ileum, the prevalence of CD4^+^ T cells (Figure 5a) was reduced in HC-fed mice compared to CTL-fed mice, though eosinophils, neutrophils, Ly6C^high^ monocytes, and total macrophages had similar prevalence between diet groups (Figure 5b-e). Colons of HC-fed mice compared to CTL-fed mice had a reduction in the prevalence of eosinophils (Figure 5g), but no differences in CD4^+^ T cells (Figure 5f), neutrophils (Figure 5h), Ly6C^high^ monocytes (Figure 5i), or total macrophages (Figure 5j). Quantities of these immune cell populations were also not different between diet groups (Supplementary Figure 4 ileum a-e; colon m-q). As we observed that small intestine and colon lengths were decreased in HC-fed mice compared to CTL-fed mice (Figure 3b-c), we also adjusted cell numbers by tissue length, which showed no differences between diet groups (Supplementary Figure 4 ileum f-j; colon r-v).

**Figure 5.**
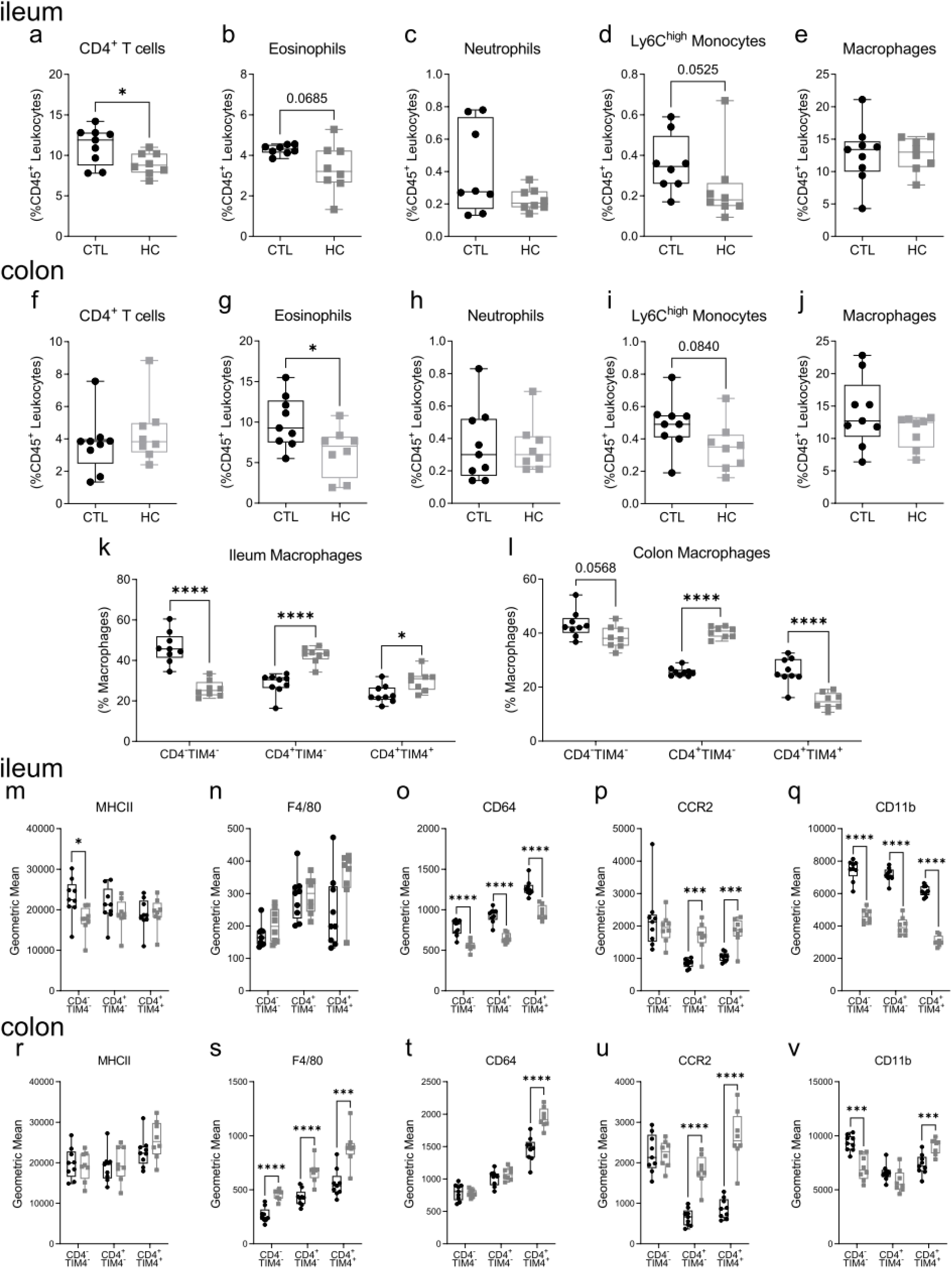
High calorie diet and excess adiposity alter maternal ileum and colon immune cell composition and macrophage phenotype in lactation. Maternal intestinal immune cell populations were assessed in lactation in ileum (**a**-**e**; **k; m-q**) and colon (**f**-**j**; **l**; **r-v**) tissues of mice fed chow (CTL) or high calorie (HC) diet. Ileum prevalence (as a proportion of CD45^+^ leukocytes) of: (**a**) CD4^+^ T cells, (**b**) eosinophils, (**c**) neutrophils, (**d**) Ly6C^high^ monocytes, (**e**) total macrophages. Colon prevalence (as a proportion of CD45^+^ leukocytes) of: (**f**) CD4^+^ T cells, (**g**) eosinophils, (**h**) neutrophils, (**i**) Ly6C^high^ monocytes, (**j**) total macrophages. Prevalence (as a proportion of total macrophages) of CD4^-^TIM4^-^, CD4^+^TIM4^-^, and CD4^+^TIM4^+^ macrophages in ileum (**k**) and colon (**l**) tissues. Ileum macrophage surface phenotype: (**m**) MHCII; (**n**) F4/80; (**o**) CD64; (**p**) CCR2; (**q**) CD11b. Colon macrophage surface phenotype: (**r**) MHCII; (**s**) F4/80; (**t**) CD64; (**u**) CCR2; (**v**) CD11b. Data are presented as box and whisker plots, minimum to maximum, with the center line at the median. Each data point indicates an individual mouse. CTL (black circles) n=8-9, HC (grey squares) n=8. Statistical significance was assessed by two-tailed Student’s t test with Welch’s correction for unequal variances or Mann-Whitney U test according to data normality. *p<0.05, ***p<0.001, ****p<0.0001.

Intestinal macrophages are either rapidly replenished from circulating Ly6C^high^ monocytes (CD4^-^TIM4^-^ and CD4^+^TIM4^-^ populations) or are locally maintained by self-proliferation (CD4^+^TIM4^+^ population)^45,46^. We next examined the prevalence of these macrophage subsets in the ileum (Figure 5k) and colon (Figure 5l), and observed tissue-specific effects of high calorie diet-induced excess adiposity. In the ileum, CD4^-^TIM4^-^ prevalence was decreased, while CD4^+^TIM4^-^ and CD4^+^TIM4^+^ populations were increased, during lactation in mice fed HC diet compared to CTL diet. The prevalence of monocyte-derived CD4^+^TIM4^-^ macrophages was likewise increased in the colons of HC-fed mice, but tissue-resident CD4^+^TIM4^+^ macrophages were decreased. These shifts in macrophage prevalence were driven by increases in numbers by tissue length of CD4^+^TIM4^-^ macrophages in both ileum and colon tissues (Supplementary Figure 4k-l, q-r). Therefore, excess adiposity influenced intestinal macrophage composition in a tissue-dependent manner, wherein the relative prevalence of self-replenishing tissue-resident macrophages are increased in the ileum but decreased in the colon, and total numbers of monocyte-derived CD4^+^TIM4^-^ macrophages are increased in both tissues.

Changes in expression of immune cell surface phenotype are often indicative of altered function. Ileum (Figure 5m-q) and colon (Figure 5r-v) macrophage surface expression was examined for markers associated with migration (CCR2, CD11b), maturity (CD64, F4/80, MHCII), and activation (CCR2, CD11b, F4/80, CD64), as well as phagocytosis (CD64) and antigen presentation (MHCII)^47–51^. Compared to CTL-fed mice, expression of CD64 and CD11b was lower on all ileal macrophage populations in HC-fed mice. In HC-fed mice compared to CTL-fed mice, there were no differences in macrophage F4/80 expression, and expression of MHCII remained lower on ileal monocyte-derived CD4^-^TIM4^-^ macrophages, whereas CCR2 was higher on monocyte-derived CD4^+^TIM4^-^ and tissue-resident CD4^+^TIM4^+^ macrophages. Colon macrophages of HC-fed mice compared to CTL-fed mice had higher expression of F4/80, and monocyte-derived CD4^+^TIM4^-^ and tissue-resident CD4^+^TIM4^+^ macrophages in the colons of HC-fed mice had higher expression of CCR2, though there were no diet-associated differences in MHCII expression across macrophage subsets. Overall, these data indicate that during lactation there were tissue-specific changes to intestinal immune cell composition and macrophage surface phenotype in mice with high calorie diet-induced excess adiposity.

### High calorie diet and excess adiposity influence maternal peripheral blood immunophenotype during lactation

As we detected an association between circulating LPS and colon paracellular permeability in mice with high calorie diet-induced excess adiposity (Figure 4j), as well as changes to the serum cytokine milieu (Figure 4k-n), we next assessed their potential systemic immunomodulatory effects, by examining immune cells in peripheral blood. We found that total leukocyte numbers (Figure 6a) and the lymphocyte to myeloid cell ratio (Figure 6b) remained similar between diet groups. HC-fed mice, compared to CTL-fed mice, had an increase in their prevalence of B cells (Figure 6c), no changes to the prevalence of CD4^+^ T cells or CD8^+^ T cells (Figure 6d), or their ratio (Figure 6e), and a decrease in NK cell prevalence (Figure 6f). The prevalence of neutrophils (Figure 6g) and total monocytes (Figure 6h) was likewise similar between HC-fed and CTL-fed mice, though there was an increase in the prevalence of Ly6C^high^ monocytes (Figure 6i). Total cell numbers of B cells, NK cells, CD4^+^ T cells, CD8^+^ T cells, neutrophils, and monocytes were not different between HC-fed and chow-fed mice (Supplementary Figure 5a-f).

**Figure 6.**
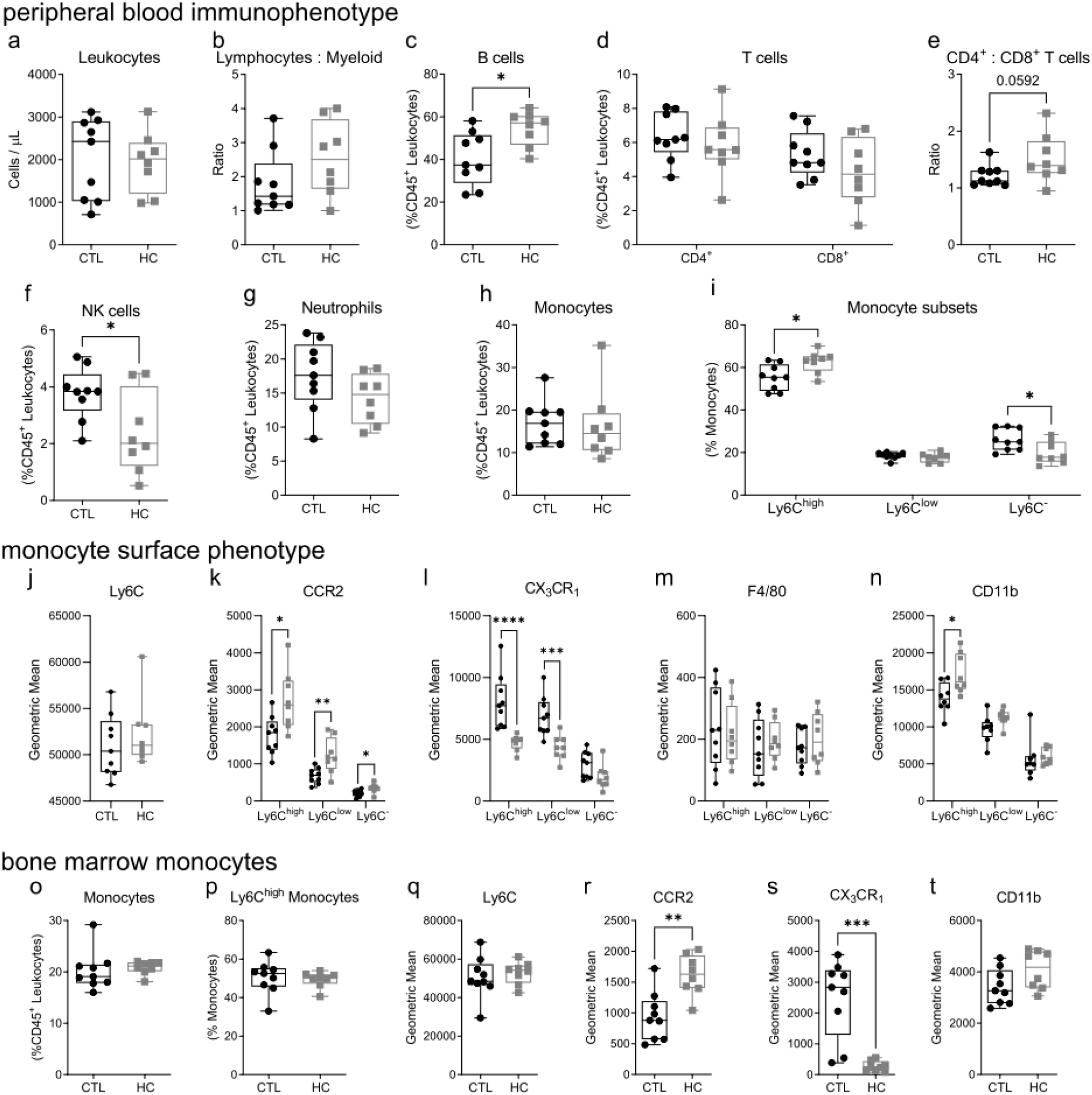
High calorie diet and excess adiposity alter maternal blood immunophenotype in lactation. Flow cytometry analysis of maternal peripheral whole blood (**a**-**n**) and bone marrow (**o**-**t**) leukocytes in lactation in mice fed chow (CTL) or high calorie (HC) diet. (**a**) CD45^+^ leukocyte cell numbers; (**b**) ratio of lymphocytes to myeloid cells. Prevalence (as a proportion of CD45^+^ leukocytes) of: (**c**) B cells, (**d**) CD4^+^ and CD8^+^ T cells. (**e**) ratio of CD4^+^ T cells to CD8^+^ T cells. Prevalence (as a proportion of CD45^+^ leukocytes) of: (**f**) NK cells, (**g**) neutrophils, (**h**) total monocytes. (**i**) prevalence of Ly6C^high^, Ly6C^low^, and Ly6C^-^ monocytes as a proportion of total monocytes. Monocyte geometric mean surface expression: (**j**) Ly6C (on Ly6C^high^ monocytes), (**k**) F4/80, (**l**) CX_3_CR_1_, (**m**) CCR2, (**n**) CD11b. (**o**) bone marrow prevalence (as a proportion of CD45^+^ leukocytes) of total monocytes. (**p**) bone marrow prevalence (as a proportion of total monocytes) of Ly6C^high^ monocytes. Bone marrow Ly6C^high^ monocyte geometric mean surface expression: (**q**) Ly6C, (**r**) CCR2, (**s**) CX_3_CR_1_, (**t**) CD11b. Data are shown as box and whisker plots, minimum to maximum, with the center line at the median. Each data point indicates an individual mouse. CTL (black circles) n=8-9, HC (grey squares) n=8. Statistical significance was assessed by two-tailed Student’s t test with Welch’s correction for unequal variances or Mann-Whitney U test according to data normality. *p<0.05, **p<0.01, ***p<0.001, ****p<0.0001.

As HC-fed mice had higher serum levels of MCP-1 (Figure 4k), a key chemokine that regulates monocyte migration and tissue infiltration, we further assessed circulating monocyte populations by examining their surface expression of CCR2 (the receptor for MCP-1), as well as Ly6C, F4/80, CX_3_CR_1_, and CD11b (Figure 6j-n). Ly6C^high^ monocytes rely on high expression of CCR2 for chemotaxis and CD11b for endothelial transmigration, and increase their expression of CX_3_CR_1_ as they mature into Ly6C^-^ monocytes, and expression of F4/80 as they mature into macrophages^47,52,53^. Blood Ly6C^high^ monocytes in CTL and HC-fed mice had similar expression of Ly6C (Figure 6j) and F4/80 (Figure 6m). Compared to CTL-fed mice, HC-fed mice Ly6C^high^ monocytes exhibited lower expression of CX_3_CR_1_ (Figure 6l), and higher expression of CCR2 (Figure 6k) as well as CD11b (Figure 6n).

We next examined monocytes within bone marrow. We observed that total monocyte and Ly6C^high^ monocyte cell numbers and prevalence were not different between the diet groups (Figure 6o-t; Supplementary Figure 5g-j). However, in HC-fed mice compared to CTL-fed mice, bone marrow Ly6C^high^ monocytes exhibited similar expression of Ly6C and CD11b but higher expression of CCR2 and lower expression of CX_3_CR_1_, (Figure 6q-t). These observations suggest that maternal excess adiposity does not have a major impact on bone marrow monocyte populations during lactation, though it may increase their tendency for CCR2-mediated migration into blood, which could contribute to the observed increase in the relative prevalence of circulating Ly6C^high^ monocytes. Overall, these data show that mice with excess adiposity had differences in peripheral blood immune cell composition and blood and bone marrow monocyte phenotype during lactation.

### High calorie diet-induced excess adiposity alters maternal metabolism and intestinal structure and permeability post-lactation

As maternal adaptations in lactation were modified by high calorie diet and excess adiposity, we next considered whether these adaptations revert to a non-pregnant state post-lactation, and examined mice up to two months after cessation of lactation. Post-lactation, mice maintained on a HC diet, compared to CTL diet, had more overall weight gain and body weight (Figure 7a; Supplementary Figure 2a-b) and excess gonadal fat (Figure 7b), but similar liver weights (Figure 7c). Though fasting insulin levels were similar between diet groups (Figure 7d), HC-fed mice also exhibited elevated fasting and non-fasting blood glucose (Figure 7e). There was a main effect of diet on oral glucose tolerance, with an increased AUC in mice with high calorie diet and excess adiposity (Figure 7f). Therefore, maintenance on the same periconceptional/perinatal diet resulted in maternal excess adiposity and hyperglycemia post-lactation.

**Figure 7.**
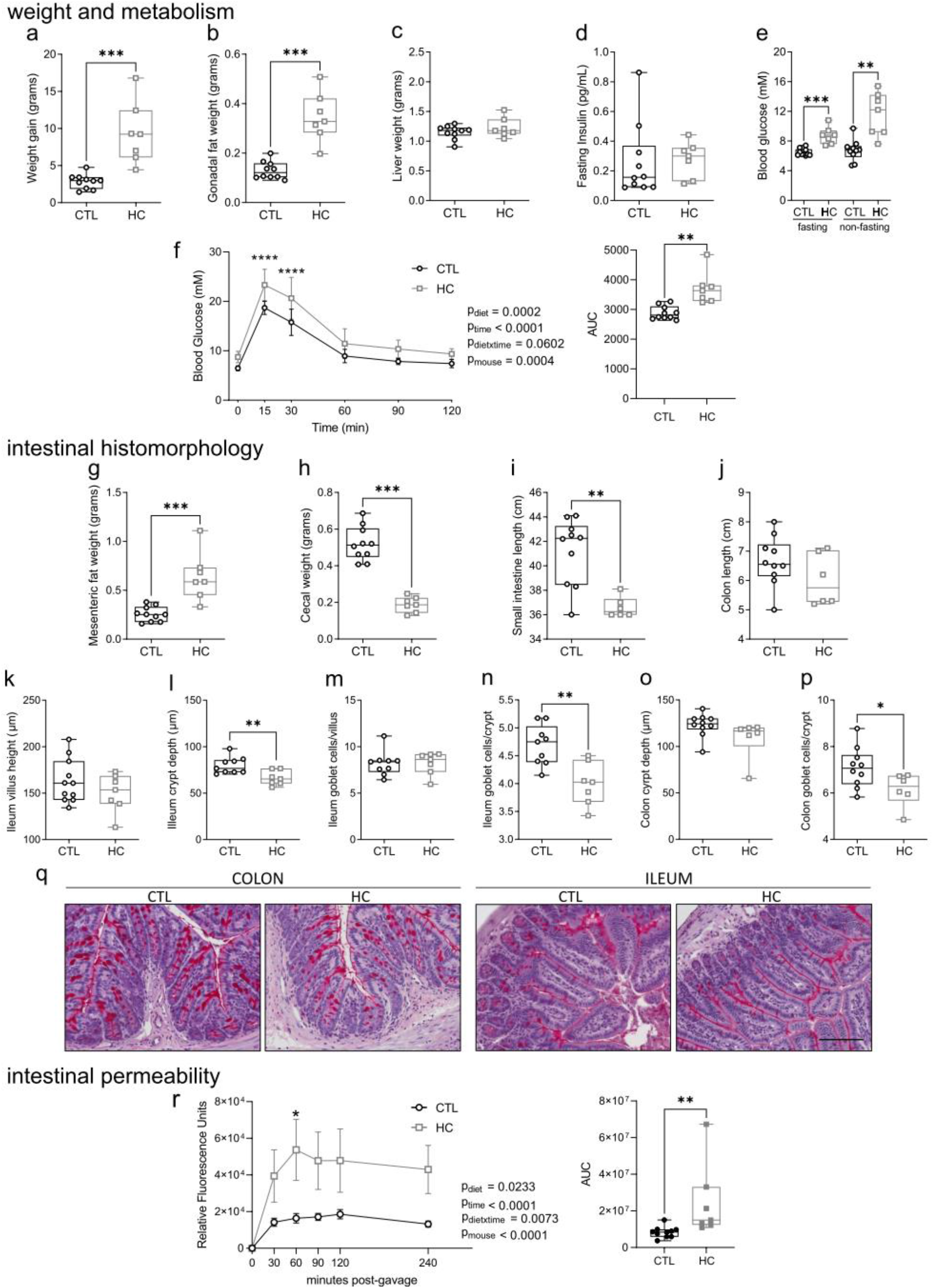
High calorie diet and excess adiposity have persistent effects on maternal metabolism and intestinal histomorphology and permeability post-lactation. Female mice allocated to chow (CTL) or high calorie (HC) diet were assessed after cessation of lactation. Maternal weight and metabolism: (**a**) maternal weight gain from start of diet allocation, (**b**) gonadal fat weight, (**c**) liver weight, (**d**) fasting blood insulin, (**e**) fasting and non-fasting blood glucose, (**f**) oral glucose tolerance test and area under the curve (AUC). Intestinal histomorphology: (**g**) mesenteric fat weight, (**h**) cecal weight, (**i**) small intestine length, (**j**) colon length, (**k**) ileum villus height, (**l**) ileum crypt depth, (**m**) ileum goblet cells per villus, (**n**) ileum goblet cells per crypt, (**o**) colon crypt depth, (**p**) colon goblet cells per crypt, (**q**) representative panels of PAS and hematoxylin staining in ileums and colons taken at 20x magnification. Scale bar: 100 µm. (**r**) *in vivo* whole-intestine permeability measured by FITC fluorescence in plasma post-gavage and area under the curve (AUC). Data in **a**-**e**, **g-p** and AUC in **f** and **r** are shown as box and whisker plots, minimum to maximum, with the center line at the median, and each data point indicates one mouse. Data in **f** are shown with a dot at the mean with error bars at ± standard deviation. Data in **r** are shown with a dot at the mean with error bars at ± standard error of the mean. CTL (black circles) n=8-10, HC (grey squares) n=4-7. Statistical significance was assessed by two-tailed Student’s t test with Welch’s correction for unequal variances or Mann-Whitney U test according to data normality (**a**-**e**, **g-p**, AUC in **f** and **r**), or two-way repeated measures ANOVA with Šidák’s post-hoc test for **f** and **r**. *p<0.05, **p<0.01, ***p<0.001,**** p<0.0001.

We next assessed if the effects of excess maternal adiposity on intestinal histomorphology and barrier function that we observed during lactation also persisted post-lactation. Mice fed HC diet compared to CTL diet had excess mesenteric fat (Figure 7g), decreased cecal weight (Figure 7h), and decreased small intestine lengths (Figure 7i), though CTL and HC-fed mice had similar colon lengths (Figure 7j). Ileum villus height was also similar between diet groups (Figure 7k), while excess adiposity was associated with reduced ileum crypt depth (Figure 7l). Goblet cell numbers were comparable in ileum villi between diet groups (Figure 7m) but decreased in ileum crypts of HC-fed mice (Figure 7n). Colon crypt depth was not different between CTL and HC-fed mice (Figure 7o), though goblet cells were reduced in colon crypts of HC-fed mice (Figure 7p). Representative PAS and hematoxylin-stained images are shown in Figure 7q. We next assessed whole-intestine permeability *in vivo* and found that there was a main effect of diet on permeability, with HC-fed mice having a higher AUC compared to CTL-fed mice (Figure 7r). Therefore, similar to our observations during lactation, maintenance on the same periconceptional/perinatal diet continued to have tissue-specific effects on intestinal structure, morphology, goblet cell numbers, and intestinal permeability post-lactation.

### High calorie diet and excess adiposity alter maternal serum cytokines and peripheral blood immunophenotype post-lactation

To assess whether systemic effects of high calorie diet-induced excess adiposity persist post-lactation, we measured serum cytokines (Figure 8a-f) and examined peripheral blood immunophenotype after the cessation of lactation (Figure 8g-s). In contrast to our observations during lactation (Figure 4k-p), we found similar serum concentrations of MCP-1 (Figure 8a), IL-6 (Figure 8b), and TNF (Figure 8c) in CTL and HC-fed mice, and HC-fed mice compared to CTL-fed mice had lower serum IL-10 (Figure 8d). As also observed during lactation, IFNγ and IL-1β concentrations were similar between diet groups (Figure 8e-f). There were no significant differences between diet groups for total leukocyte numbers (Figure 8g), the lymphocyte to myeloid cell ratio (Figure 8h), the prevalence of B cells (Figure 8i), CD4^+^ or CD8^+^ T cells (Figure 8j), their ratio (Figure 8k), or the prevalence of NK cells (Figure 8l). Numbers of these immune cell populations were also similar between diet groups (Supplementary Figure 6a-d). In contrast, HC-fed mice had a reduced prevalence of neutrophils (Figure 8m), though neutrophil numbers were similar between diet groups (Supplementary Figure 6e). While the prevalence of total monocytes and monocyte subset composition was similar in HC and CTL-fed mice (Figure 8n-o), total numbers of monocytes were significantly increased in HC-fed mice (Supplementary Figure 6f). Further characterization of monocyte surface phenotype showed that Ly6C^high^ monocytes had increased expression of Ly6C (Figure 8p), but expression of CCR2, F4/80, and CD11b was not different between diet groups for any monocyte subset (Figure 8q-s). Therefore, HC diet and excess adiposity influenced peripheral blood myeloid cells post-lactation.

**Figure 8.**
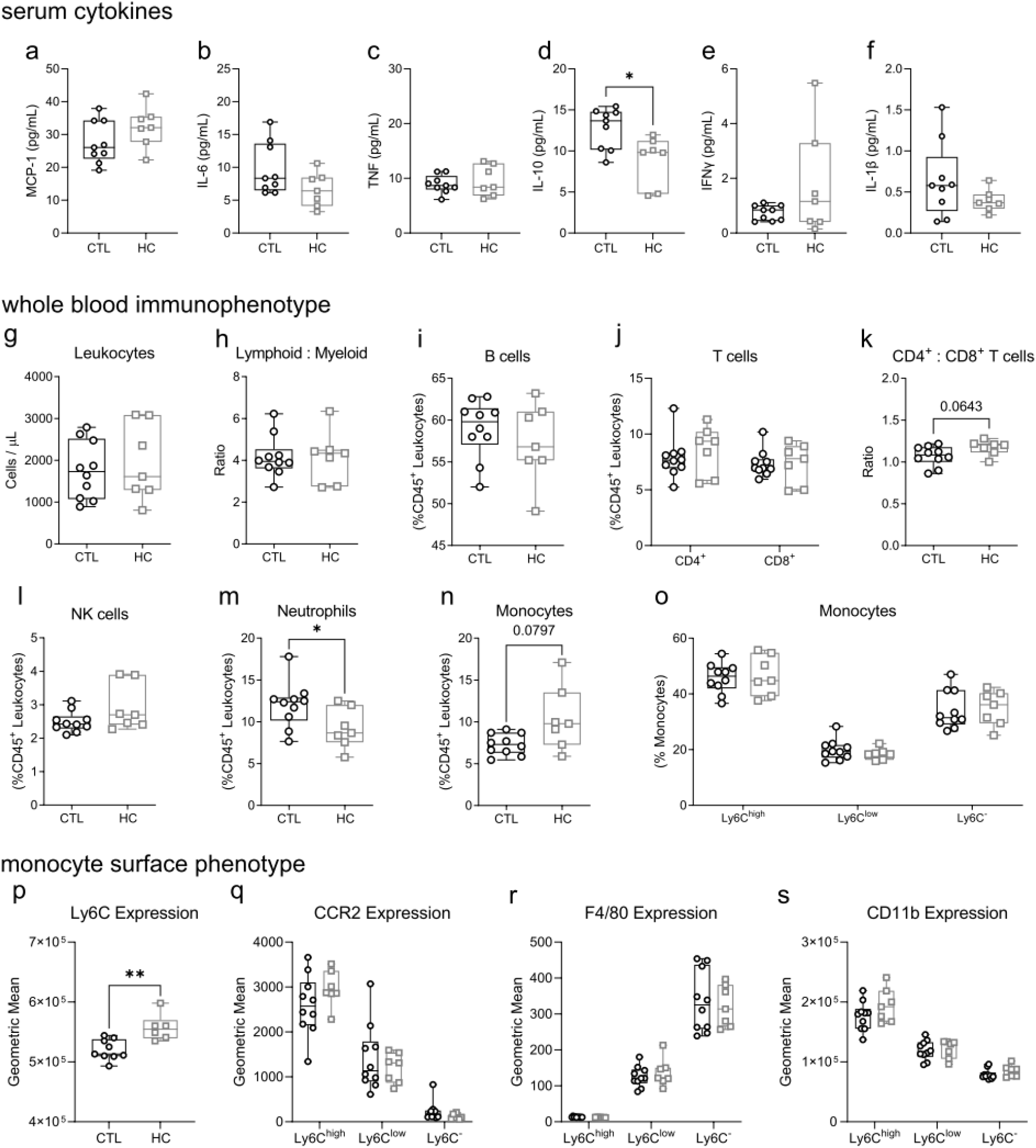
High calorie diet and excess adiposity influence maternal systemic inflammation and blood immunophenotype post-lactation. Assessments of maternal circulating immunomodulatory factors and immune cell composition post-lactation in mice fed chow (CTL) or high calorie (HC) diet. Circulating cytokines: (**a**) MCP-1, (**b**) IL-6, (**c**) TNF, (**d**) IL-10, (**e**) IFNγ, (**f**) IL-1β. (**g**) CD45^+^ leukocyte cell numbers. (**h**) ratio of lymphocytes to myeloid cells. Prevalence (as a proportion of CD45^+^ leukocytes) of: (**i**) B cells, (**j**) CD4^+^ and CD8^+^ T cells. (**k**) ratio of CD4^+^ T cells to CD8^+^ T cells. Prevalence (as a proportion of CD45^+^ leukocytes) of: (**l**) NK cells, (**m**) neutrophils, (**n**) total monocytes. (**o**) prevalence of Ly6C^high^, Ly6C^low^, and Ly6C^-^ monocytes as a proportion of total monocytes. Monocyte geometric mean surface expression: (**p**) Ly6C on Ly6C^high^ monocytes; (**q**) CCR2; (**r**) F4/80; (**s**) CD11b. Data are shown as box and whisker plots, minimum to maximum, with the center line at the median, and each data point indicates an individual mouse. CTL (black circles) n=10, HC (grey squares) n=7. Statistical significance was assessed by two-tailed Student’s t test with Welch’s correction for unequal variances or Mann-Whitney U test according to data normality. **p<0.01, *p<0.05.

### Lactation and diet modify maternal metabolism, intestinal permeability, and peripheral blood immunophenotype

To provide further insights into the potential post-lactation resolution of maternal adaptations observed during lactation, and effects of excess adiposity, we directly compared our data collected in lactation and post-lactation on maternal glucose tolerance (Figure 9a), intestinal permeability (Figure 9b-d), and peripheral blood immunophenotype (Figure 9e-k). There was a was main effect of time on glucose tolerance, with HC-fed mice post-lactation having significantly higher AUC than HC-fed mice in lactation, as well as higher AUC compared to CTL-fed mice post-lactation (Figure 9a). In contrast, CTL-fed mice had similar glucose tolerance in lactation and post-lactation. Intestinal permeability, measured by *in vivo* assay and AUC, was modified by a main effect of maternal diet, and was particularly enhanced post-lactation in HC-fed mice compared to CTL-fed mice (Figure 9b). Data were also compared to assessments in never-pregnant virgin mice. In mice fed CTL diet in lactation and post-lactation (Figure 9c), there was a main effect of lactation on intestinal permeability, with never-pregnant and post-lactation mice having more similar permeability than mice in lactation (Figure 9c). In mice fed HC diet, intestinal permeability was significantly increased in both lactation and post-lactation compared to never-pregnant mice.

**Figure 9.**
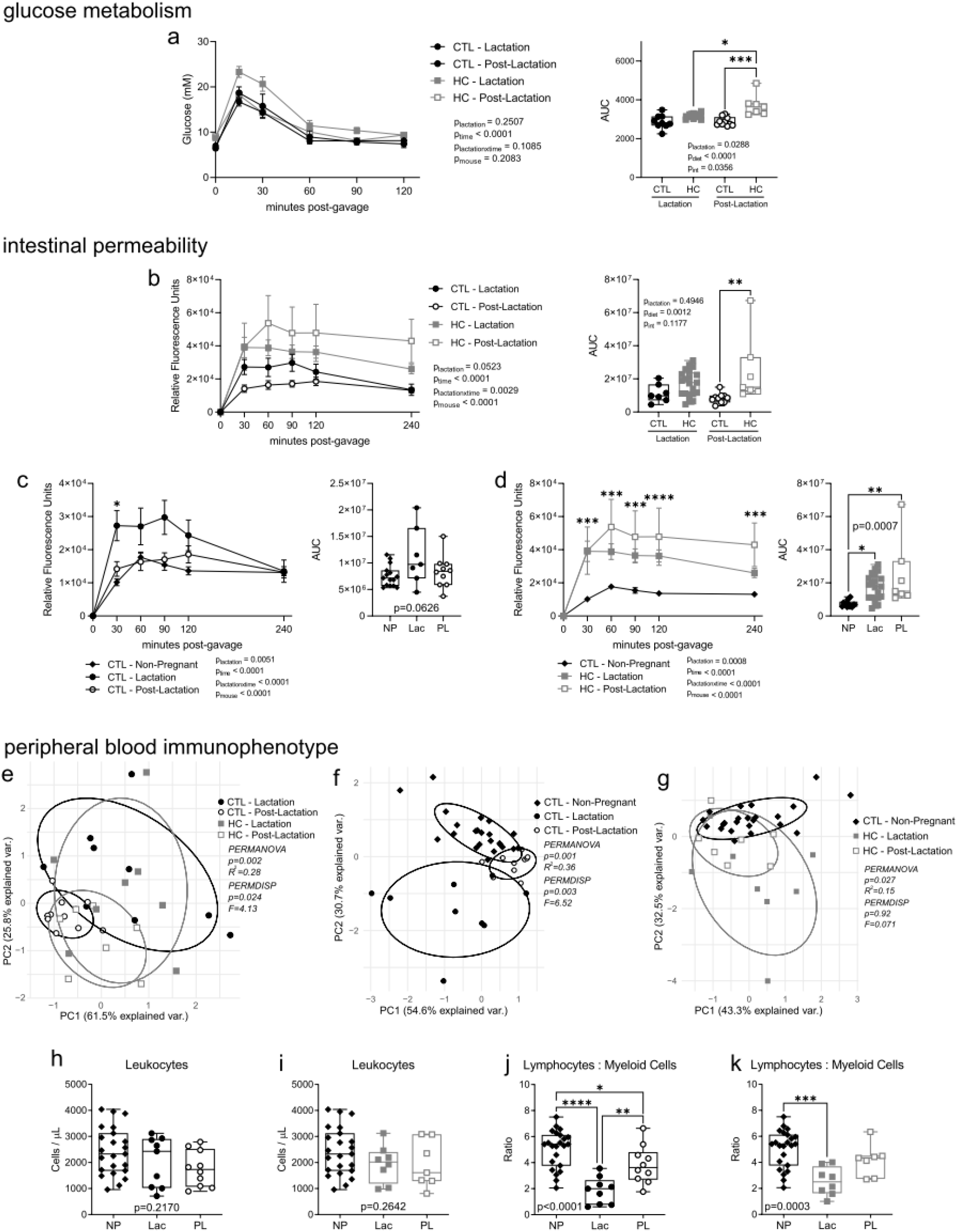
Comparison of maternal glucose metabolism, intestinal permeability and peripheral immunophenotype during lactation and post-lactation. Comparisons between maternal assessments in lactation (Lac; from Figures 2, 4, and 6) and post-lactation (PL; from Figures 7 and 8) and age-matched never/non-pregnant virgin female mice (NP). (**a**) oral glucose tolerance test and area under the curve (AUC). (**b-d**) *in vivo* whole-intestine permeability measured by FITC fluorescence in plasma post-gavage and AUC. (**e-g**) PCA of peripheral blood immunophenotype. (**h-i**) total blood CD45^+^ leukocyte numbers. (**j-k**) ratio of blood lymphocytes to myeloid cells. Data in **a** are shown with a dot at the mean with error bars at ± standard deviation. Data in **b-d** are shown with a dot at the mean with error bars at ± standard error of the mean. Data for AUC in **a-d** and data in **h-k** are shown as box and whisker plots, minimum to maximum, with the center line at the median, and each data point indicates an individual mouse. Data in **e-g** from PCA are plotted on PC1 and PC2, and each data point indicates an individual mouse. Mice were allocated to chow (CTL) or high calorie (HC) diet. CTL Lac (black circles) n=9; CTL PL (open black circles) n=10; HC Lac (grey squares) n=8 or n=21 (for **b** and **d**); HC PL (open grey squares) n=7; CTL NP (black diamonds) n=15 (for **c-d**) or n=23 (for **e-k**). Statistical significance for line graphs in **a-d** was assessed by repeated measures ANOVA with Šidák’s correction for multiple comparisons (post-hoc comparisons not shown for **b** and comparisons shown in **c** and **d** on line graphs are between CTL-Lac and NP, and HC-PL and NP, respectively), for AUC in **a** and **b** by two-way ANOVA with Tukey’s multiple comparisons correction, for AUC in **c-d** and data in **h-k** by one-way ANOVA with Tukey’s correction (parametric) or Kruskal-Wallis with Dunn’s correction (non-parametric), and for **e-g** using the R functions *adionis2*, *permutest* and *betadisper*. *p<0.05, ***p<0.01,*** p<0.001, ****p<0.0001.

Changes to blood immunophenotype were initially considered on a whole-population basis (Figure 9e-g). Immune cell composition changed between lactation and post-lactation, with less similarity (indicated by less population overlap) between lactation and post-lactation immunophenotype in CTL-fed mice compared to lactation and post-lactation immunophenotype in HC-fed mice with excess adiposity (Figure 9e). Differences in immunophenotype between lactation and post-lactation were also more apparent in CTL-fed mice than HC-fed mice when compared to never-pregnant CTL-fed female mice (Figures 9f-g). Effects of diet, as well as lactation and post-lactation resolution, were in addition considered in context of individual immune cell populations (Figure 9h-k; Supplementary Figure 7). While total leukocyte numbers were similar in never-pregnant mice and mice in lactation and post-lactation (Figure 9h-i), the ratio of lymphocytes to myeloid cells significantly decreased in lactation in both CTL and HC-fed mice (Figure 9j-k). This ratio also remained lower post-lactation in CTL-fed mice but not HC-fed mice with excess adiposity, compared to never-pregnant mice. Differences in other maternal immune cell adaptations during lactation and their resolution post-lactation were apparent between CTL and HC-fed mice with excess adiposity (Supplementary Figure 7). Intestinal immune cell composition also significantly changed between CTL-fed never-pregnant mice and during lactation, with similar shifts in ileum and colon tissues, which were modified by high calorie diet-induced excess adiposity (Supplementary Figure 8). Therefore, maternal glucose metabolism, intestinal permeability, and immunophenotype were modified by adaptations in lactation, and these adaptations, as well as their persistence and resolution, were influenced by maternal diet and excess adiposity.

## Discussion

With the rising global incidence of excess weight gain in the periconceptual/perinatal period^54^, and associated postpartum weight retention^8^, which predict short and long-term negative health outcomes^5,6^, it is important to understand how excess adiposity modifies maternal adaptations, particularly in the rarely considered early postpartum period of lactation. In this study, as summarized in Figure 10, we found that maternal metabolic, gastrointestinal, and immunological adaptations during lactation in mice are modified by high calorie diet-induced excess adiposity, and that many of the observed changes persisted two months post-lactation.

**Figure 10.**
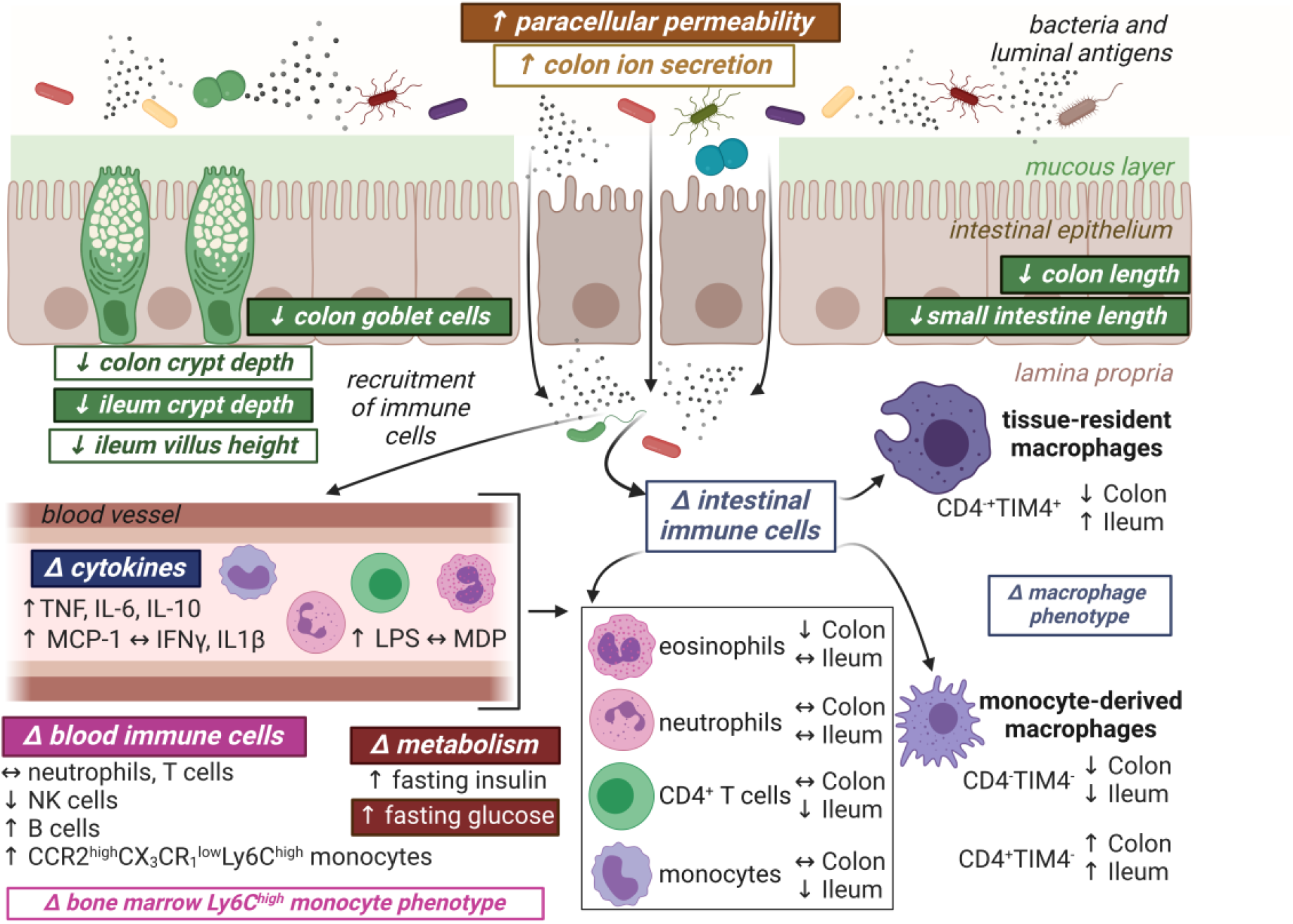
Summary of effects of high calorie diet and excess adiposity on maternal adaptations in lactation. Effects of high calorie diet and excess adiposity also observed up to two months post-lactation are indicated by shaded text boxes.

The pathophysiology of excess adiposity and its accompanying immunometabolic dysregulation are associated with impaired intestinal barrier function in non-pregnant female mice^22^, and in pregnancy^14^. In this study we extended those observations, showing that in the early postpartum period in lactation, and up to two months post-lactation, mice with excess adiposity have increased intestinal permeability. Changes to intestinal histomorphology and local immune cells in mice with excess adiposity likely contributed to impaired intestinal barrier function, and *vice versa*. In particular, we found that maternal ileum crypt depth and ileum villus height were decreased in mice with excess adiposity during lactation, consistent with our previous observations in non-pregnant female mice, and pregnant mice at mid- and late-gestation^14^. These morphological changes may be indicative of altered intestinal stem cell proliferation, as has previously been reported in non-pregnant female mice with diet-induced obesity^24^. Goblet cells support formation of the mucous barrier overlying the intestinal epithelium^20^, through secretion of mucins including *Muc2*, which has decreased expression with excess adiposity in female non-pregnant^24^ and pregnant mice at mid- and late-gestation^29,30^. In lactation, and post-lactation, we found that there was a reduction in goblet cells per crypt in the colon, as has also been reported in non-pregnant female mice^14,24^ and pregnant mice at mid-gestation^14^ with diet-induced excess adiposity. In addition, we have previously found increased ileal gene expression of *Cd4* as well as *F4/80* (commonly expressed by tissue macrophages) in late gestation in mice fed a high fat high calorie diet^28^. In this study we specifically identified intestinal immune cells and found tissue-specific differences in composition. Moreover, in mice with excess adiposity during lactation, the prevalence and numbers of monocyte-derived CD4^+^TIM4^-^ macrophages, with altered surface phenotype, were increased in both the ileum and colon. CD4^+^TIM4^-^ macrophages localize proximal to the intestinal epithelium to support its maintenance^46^, so this increase may be reflective of requirements to support intestinal barrier function in mice with excess adiposity during lactation. These observations differ from those we have previously reported in non-pregnant mice with excess adiposity^22^, but this was not unexpected, as we also found in this study that intestinal macrophage population dynamics are altered in lactation, independent of maternal diet. Therefore, impaired intestinal barrier function, and changes to local cellular structure and macrophage phenotype, accompany high calorie diet-induced excess adiposity in lactation.

Intestinal permeability may lead to dissemination of microbial components into systemic circulation, with modulation of peripheral inflammation and immune cell composition^19^. We observed elevated levels of the bacterial cell wall component LPS, as well as the pro-inflammatory cytokines IL-6 and TNF, in circulation of mice with excess adiposity in lactation, indicative of systemic inflammation. We also found elevated MCP-1, which regulates migration of monocytes, whether from bone marrow into circulation, or from circulation into tissues^47^. Monocytes and monocyte-derived tissue macrophages in non-pregnant mice, and their production of proinflammatory cytokines, contribute to the pathophysiology of excess adiposity and accompanying metabolic and immunological changes^27,55,56^. In this study, irrespective of maternal diet, we observed that the expansion of peripheral blood neutrophils and reduction of B cells that occurs in pregnancy^57,58^ persists into lactation. As has been reported in dairy cows^59^, we also observed an increase in circulating monocytes in lactation. In mice with excess adiposity, Ly6C^high^ monocytes in both bone marrow and blood had increased surface expression of the MCP-1 receptor CCR2, and lower surface expression of CX_3_CR_1_, which mediates retention of monocytes in the bone marrow. This surface phenotype is also observed on circulating Ly6C^high^ monocytes in non-pregnant female mice with diet-induced obesity^27^. However, while the relative prevalence of Ly6C^high^ monocytes increased in circulation of mice with excess adiposity in lactation, total cell numbers in blood and bone marrow did not increase. This dichotomy could reflect the specific time point of assessment in lactation, as we previously found that diet-induced obesity increased circulating Ly6C^high^ monocytes at mid-but not late-gestation^14^, and may also be a consequence of the rapid egress of monocytes into tissues including mammary glands^60^. Pregravid obesity in humans has been reported to attenuate pregnancy-associated metabolic, transcriptional and epigenetic adaptations within monocytes^37^. Combined, these observations prompt future investigations of maternal monocytes to determine if excess adiposity in the periconceptual/perinatal period leads to monocyte epigenetic reprogramming that promotes a long-term inflammatory and obesogenic phenotype, as has been reported in male mice with diet-induced obesity^61^.

The postpartum resolution of maternal adaptations after pregnancy and lactation is not an instantaneous process. In this study, we observed in mice without excess adiposity that within two months post-lactation, not all peripheral blood immune cell populations had returned to pre-pregnancy levels. We found in contrast that whole-intestine permeability decreased in absence of excess adiposity between lactation and post-lactation, and was similar between never-pregnant female mice and mice two months post-lactation, suggesting that barrier function returns to pre-pregnancy levels after the cessation of lactation. Yet, it has been reported that enhanced intestinal nutrient transport capacity remains for up to 70 days postpartum, higher small intestine villi remain for up to 200 days, and small intestine length remains longer for up to 300 days^15^. Thus, certain pregnancy/lactation adaptations even in the absence of excess adiposity could remain for a significant part of the lifespan. We in addition found post-lactation that mice with high calorie diet-induced excess adiposity continued to have increased whole-intestine permeability and altered intestinal structure, as well as differences in peripheral immunomodulatory factors and immunophenotype that were distinct from those observed during lactation. Therefore, the post-lactation resolution of maternal adaptations was impaired in mice with excess adiposity. Even independent of postpartum diet, excess adiposity during gestation in mice has been reported to increase risk of obesity up to nine months post-pregnancy^62^. Future investigations should consider the periconceptual/perinatal timing of weight gain, and whether this further modifies effects of excess adiposity observed during lactation in this study, as a higher risk of postpartum obesity has been reported in women with excess weight gain in the first trimester compared to the second or third trimester^63^. These data also imply that parity may be an important consideration in development of mouse models for conditions that are more prevalent in women.

The focus of this study was on maternal effects of lactation and excess adiposity due to a high calorie diet, but maternal excess adiposity in pregnancy is known to impact offspring metabolism, immunity, and neurodevelopment^64–67^. Changes to intestinal physiology and peripheral and intestinal immune cell composition that we observed in mice with excess adiposity during lactation likely also influence the composition of breast milk, which could have long-term effects on offspring development and health^68,69^. Overall, this study offered new insights into the vulnerability of maternal intestinal, metabolic, and immune adaptations in lactation to excess adiposity, and emphasizes the need for more studies on maternal postpartum health to develop effective interventions to improve outcomes of pregnancy and lactation, as well as the trajectory of lifelong maternal health.

## Author Contributions

J.A.B. contributed study concept and design, data acquisition, data analysis, interpretation of data, writing – initial manuscript, writing – editing and revision

T.A.R. contributed study concept and design, data acquisition, data analysis, interpretation of data, writing – editing and revision

E.Y. contributed data acquisition, data analysis, interpretation of data, writing - editing and revision

B.K.E.K. contributed data acquisition

X.W. contributed data acquisition, data analysis, interpretation of data

C.Q. contributed data acquisition

B.C. contributed data acquisition

E.F.V. contributed supervision, provision of resources, writing – editing and revision

D.M.E.B. contributed study concept and design, supervision, design of project, provision of resources, writing – editing and revision

D.M.S. contributed study concept and design, funding, supervision, design of project, provision of resources, writing - editing and revision

## Acknowledgements

The authors would like to thank the McMaster University Central Animal Facility staff and veterinarians for their care of the animals in this study, Hong Liang and the McMaster Immunology Research Centre flow cytometry facility, Erin Demask and Tina Walker for provision of media, John Grainger for guidance on intestinal tissue processing and flow cytometry protocol design, and Katherine Kennedy and Patrycja Jazwiec for their assistance with tissue collections. Figures 1 and 10 were made using BioRender.com.

---

**Supplementary Figure 1.**
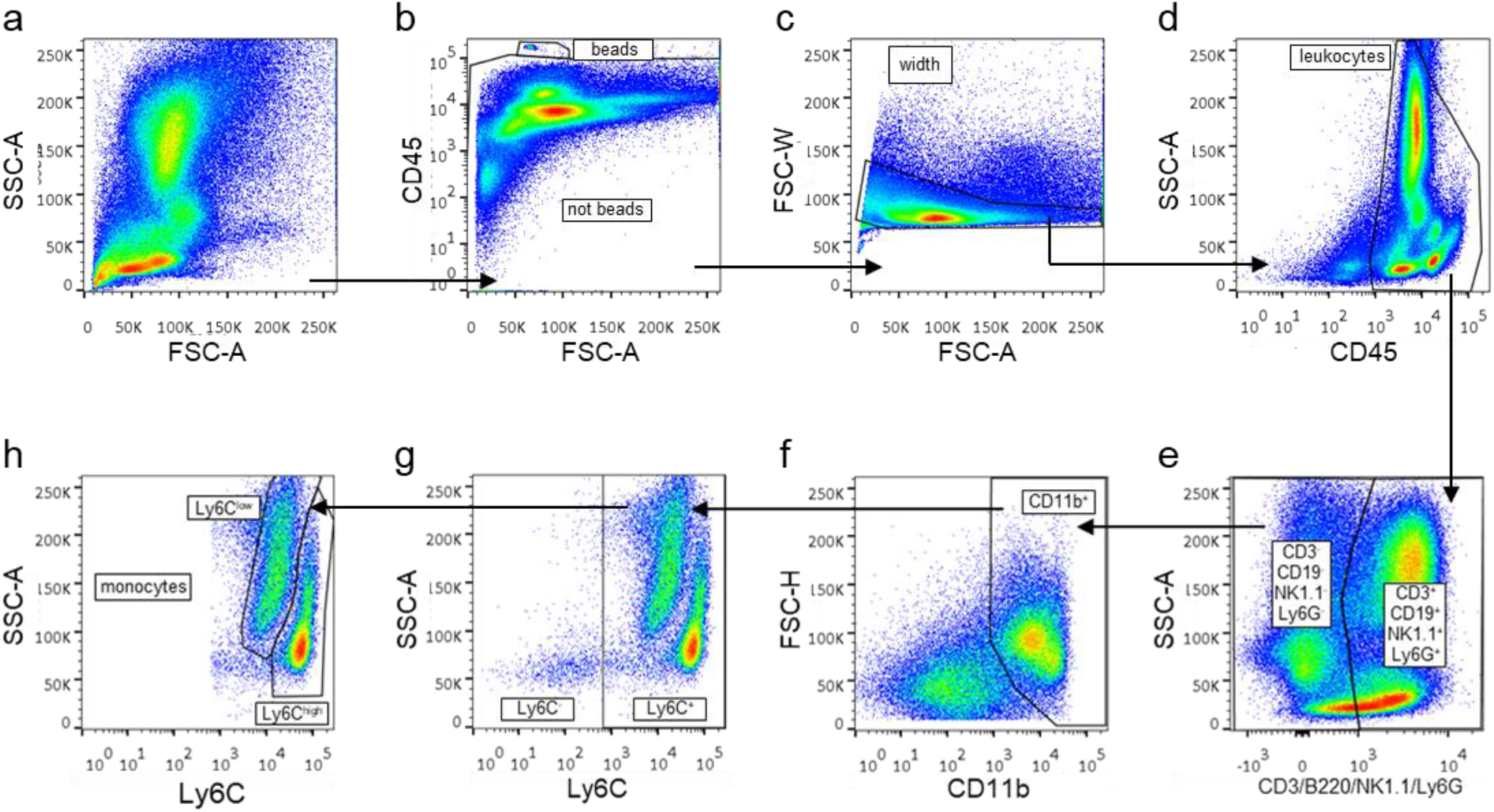
Gating strategy to identify bone marrow monocytes by surface staining and flow cytometry. A representative sample during lactation from a mouse fed the chow diet is shown. (**a**) captured events. (**b)** absolute count beads (beads) are separated from collected events (not beads) as CD45^high^ events. (**c**) a width gate is applied to exclude aggregates. (**d**) CD45^+^ leukocytes are gated. (**e**) a CD3/CD19/NK1.1/Ly6G dump gate is used to remove T cells (CD3^+^), B cells (CD19^+^), NK cells (NK1.1^+^), and neutrophils (Ly6G^+^). (**f**) CD45^+^CD3^-^CD19^-^NK1.1^-^Ly6G^-^ cells are gated according to their expression of CD11b to identify remaining myeloid cells (CD11b^+^). (**g**) CD45^+^CD3^-^CD19^-^NK1.1^-^Ly6G^-^CD11b^+^ cells are gated according to their expression of Ly6C to identify Ly6C^+^ and Ly6C^-^ populations (CD45^+^CD3^-^CD19^-^NK1.1^-^Ly6G^-^CD11b^+^Ly6C^+/-^). (**h**) Ly6C^+^ cells are divided into Ly6C^high^ and Ly6C^low^ monocytes.

**Supplementary Figure 2.**
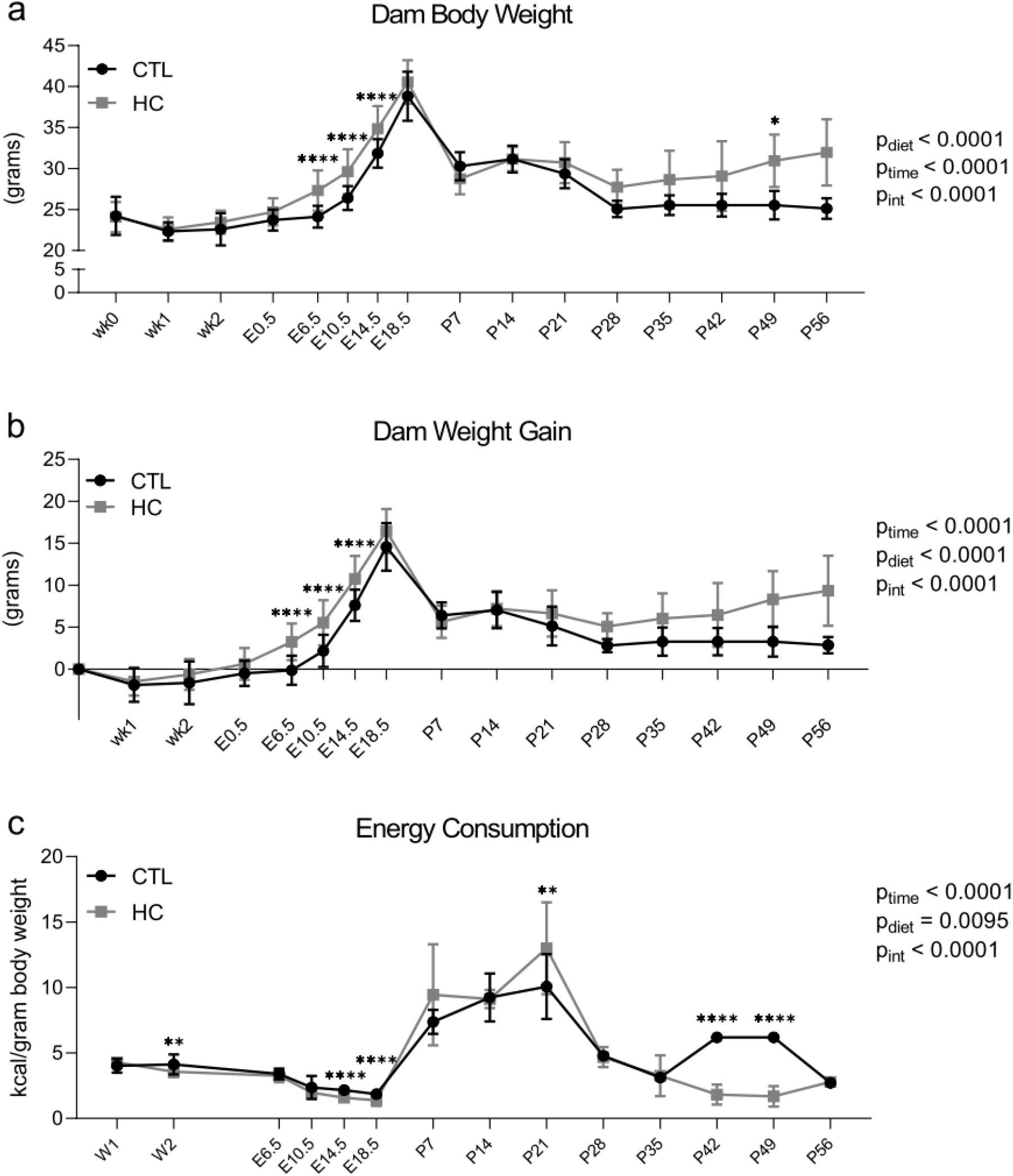
Time course of maternal body weight, weight gain, and energy consumption. Female mice were allocated to chow (CTL) or high calorie (HC) diet. From start of diet allocation pre-pregnancy (W1, W2) through pregnancy (E6.5 to E18.5), lactation (P7 to P21), and post-lactation (P28 to P56): (**a**) maternal body weight, (**b**) maternal weight gain, (**c**) maternal energy consumption. Data are shown with a dot at the group mean ± standard deviation. CTL n=34 (to P21) or n=10 (to P56), HC n=39 (to P21) or n=7 (to P56). Statistical significance was assessed between diet groups at each time point by repeated measures mixed effects model, with Šidák’s multiple comparisons post-hoc test. **p<0.01, ****p<0.0001.

**Supplementary Figure 3.**
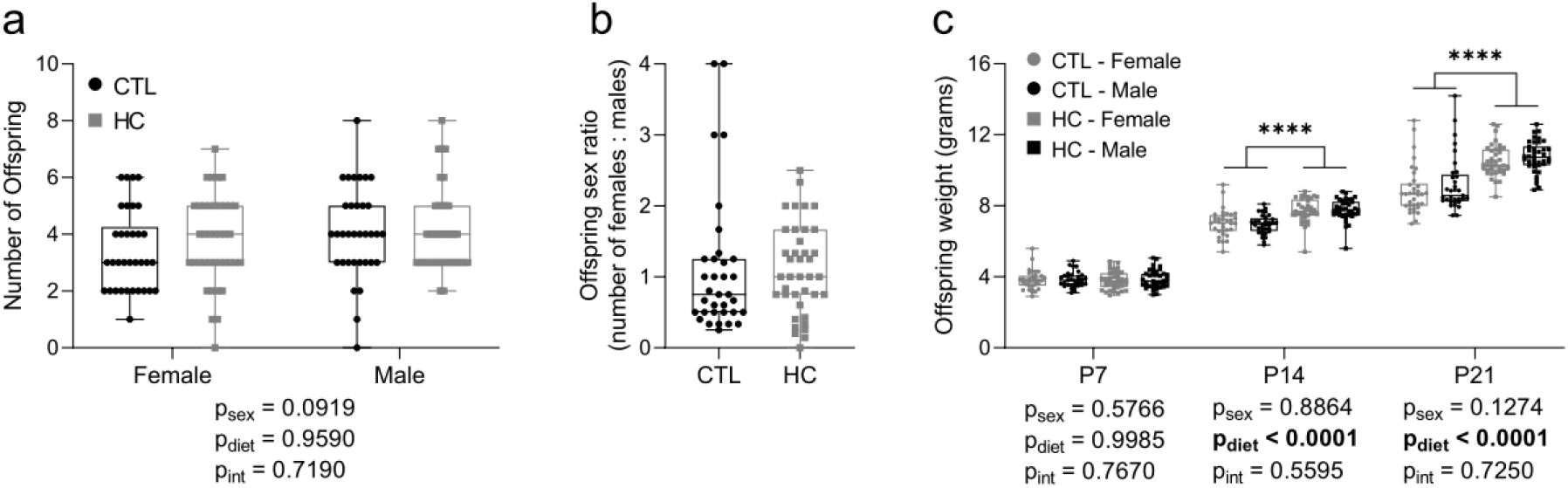
Maternal high calorie diet and excess adiposity do not significantly alter offspring numbers or sex ratio but increase offspring weight during lactation. Offspring assessments from female mice allocated to chow (CTL) or high calorie (HC) diet. (**a**) number of female and male offspring at P7. (**b**) offspring female to male sex ratio at P7. (**c**) weights of female and male offspring at P7, P14 and P21. Data are shown as box and whisker plots, minimum to maximum, with the center line at the median. Each data point represents one dam (i.e., offspring measures were averaged within a litter per dam). For **a**, CTL female n=34, male n=33, HC female n=38, male n=39. For **b**, CTL n=34, HC n=39. For **c**, CTL female n=34, male n=33, HC female n=37-8, male n=38-39. Statistical significance was assessed in **a** and **c** by two-way ANOVA with Tukey’s post-hoc test, in **b** by Mann-Whitney U test. Also see Supplementary Table 2. ****p<0.0001.

**Supplementary Figure 4.**
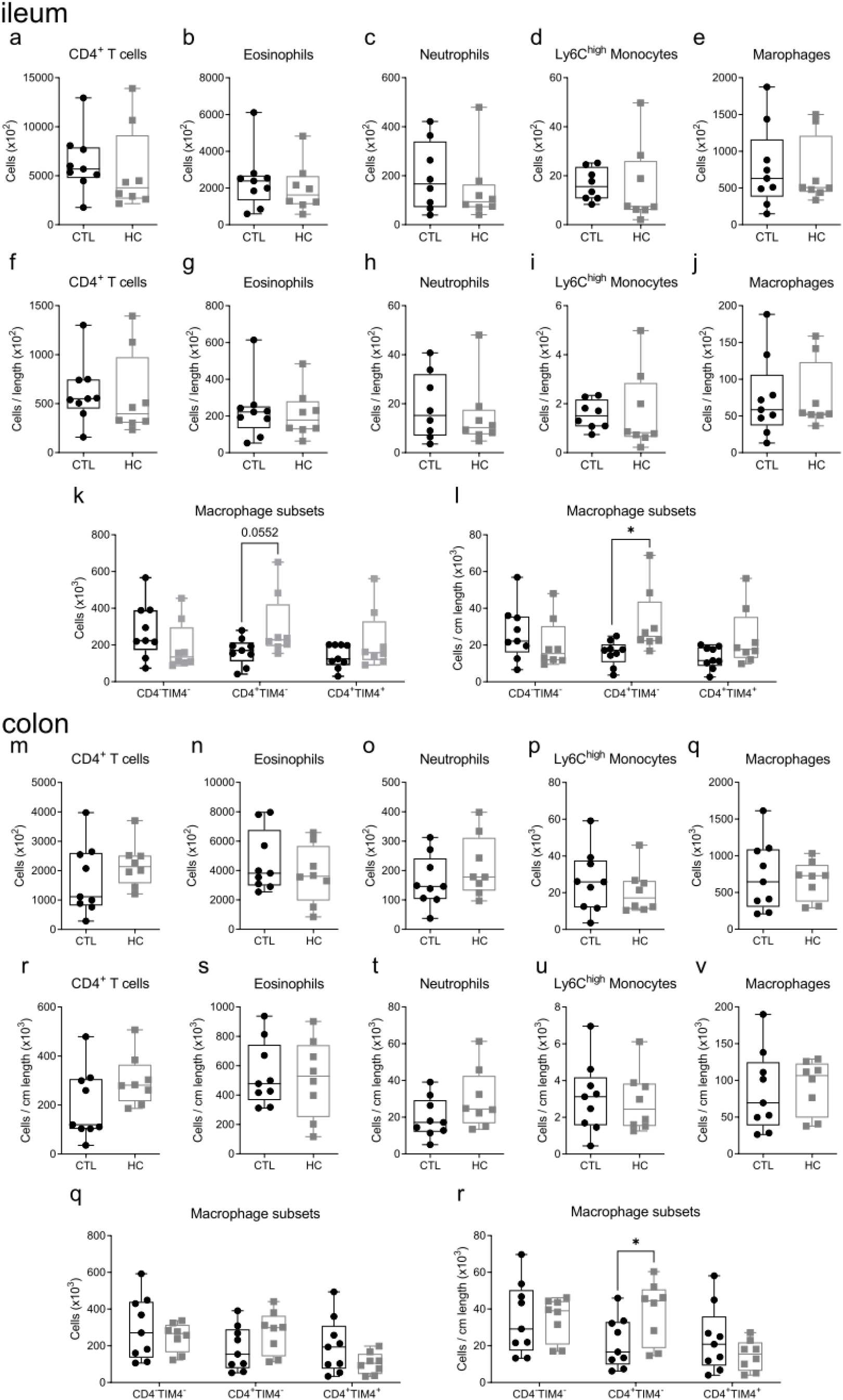
Effects of high calorie diet and excess adiposity on numbers of maternal ileum and colon immune cells in lactation. Maternal intestinal immune cell populations in the ileum (**a**-**l**) and colon (**m**-**x**) of mice during lactation fed chow (CTL) or high calorie (HC) diet. Ileum cell numbers of: (**a**) CD4^+^ T cells, (**b**) eosinophils, (**c**) neutrophils, (**d**) Ly6C^high^ monocytes, (**e**) total macrophages. Ileum cell numbers by tissue length of: (**f**) CD4^+^ T cells, (**g**) eosinophils, (**h**) neutrophils, (**i**) Ly6C^high^ monocytes, (**j**) total macrophages. Ileum CD4^-^TIM4^-^, CD4^+^TIM4^-^, and CD4^+^TIM4^+^ macrophages: (**k**) cell numbers, (**l**) cell numbers by tissue length. Colon cell numbers of: (**m**) CD4^+^ T cells, (**n**) eosinophils, (**o**) neutrophils, (**p**) Ly6C^high^ monocytes, (**q**) total macrophages. Colon cell numbers by tissue length of: (**r**) CD4^+^ T cells, (**s**) eosinophils, (**t**) neutrophils, (**u**) Ly6C^high^ monocytes, (**v**) total macrophages. Colon CD4^-^TIM4^-^, CD4^+^TIM4^-^, and CD4^+^TIM4^+^ macrophages: (**w**) cell numbers; (**x**) cell numbers by tissue length. Data are presented as box and whisker plots, minimum to maximum, with the center line at the median, and each data point indicates an individual mouse. CTL (black circles) n=9, HC (grey squares) n=8. Statistical significance was assessed by two-tailed Student’s t test with Welch’s correction for unequal variances or Mann-Whitney U test according to data normality. *p<0.05.

**Supplementary Figure 5.**
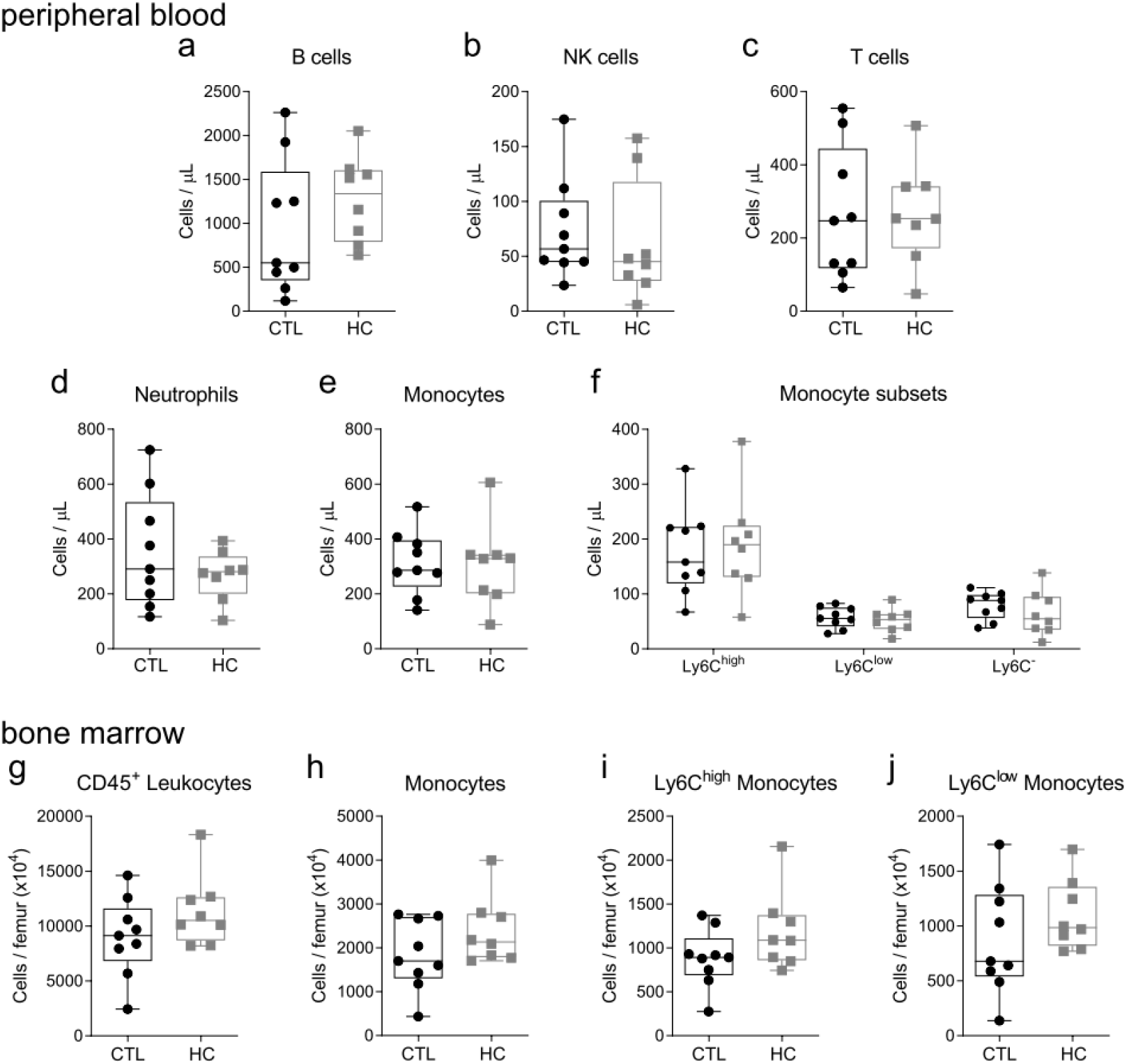
Effects of high calorie diet and excess adiposity on maternal peripheral blood and bone marrow immunophenotype cell numbers during lactation. Maternal peripheral blood (**a**-**f**) and bone marrow (**g**-**j**) leukocytes of mice fed chow (CTL) or high calorie (HC) diet. Peripheral blood cell numbers of: (**a**) B cells, (**b**) NK cells, (**c**) T cells, (**d**) neutrophils, (**e**) total monocytes, (**f**) Ly6C^high^, Ly6C^low^, and Ly6C^-^ monocytes. Bone marrow cell numbers of: (**g**) total CD45^+^ leukocytes, (**h**) total monocytes, (**i**) Ly6C^high^ monocytes, (**j**) Ly6C^low^ monocytes. Data are presented as box and whisker plots, minimum to maximum, with the center line at the median, and each data point indicates an individual mouse. CTL (black circles) n=9, HC (grey squares) n=8. Statistical significance was assessed by two-tailed Student’s t test with Welch’s correction for unequal variances or Mann-Whitney U test according to data normality.

**Supplementary Figure 6.**
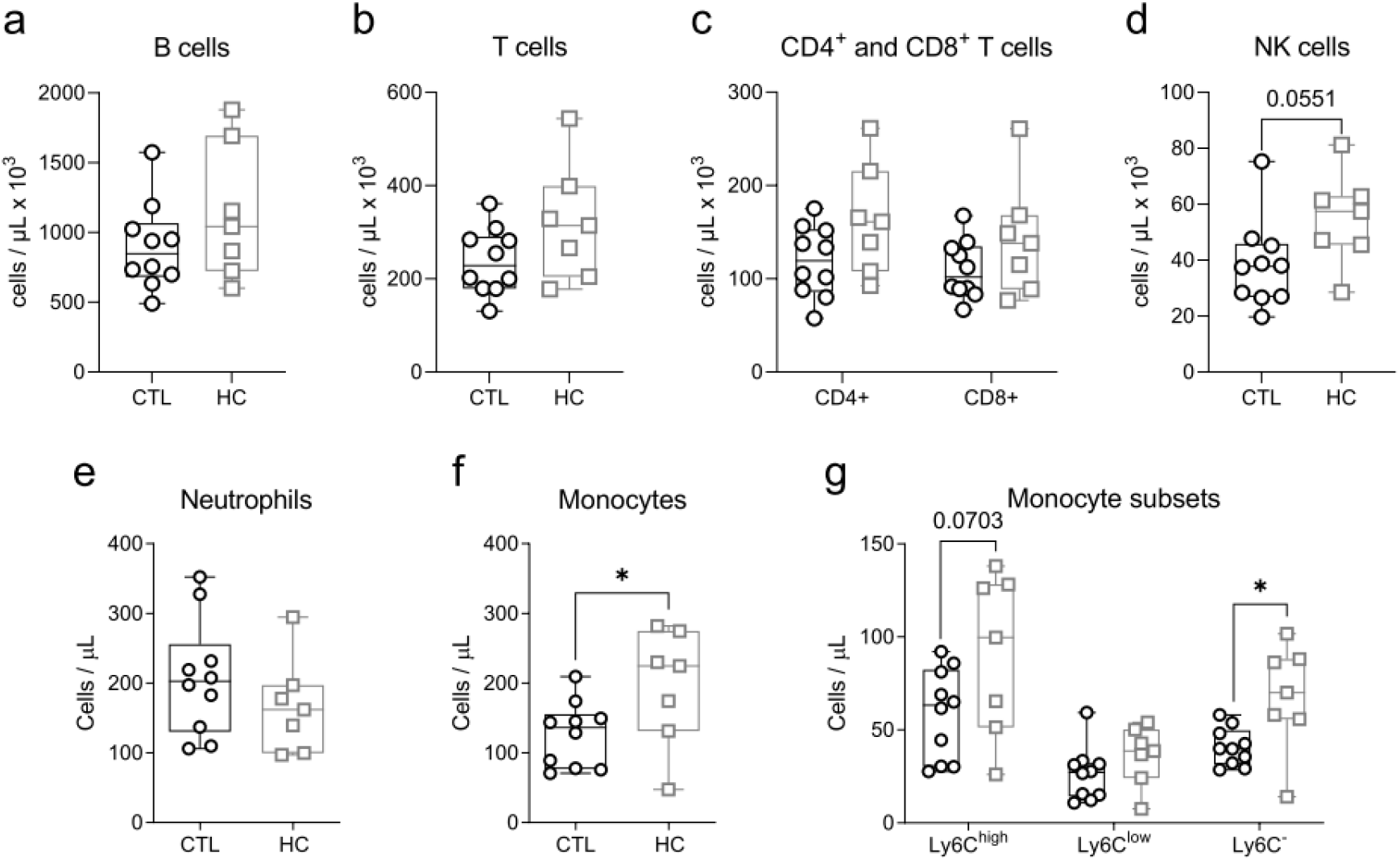
Effects of high calorie diet and excess adiposity on maternal peripheral blood immunophenotype cell numbers post-lactation. Maternal peripheral blood immunophenotype cell counts of mice fed chow (CTL) or high calorie (HC) diet. Numbers of: (**a**) B cells, (**b**) T cells, (**c**) CD4^+^ and CD8^+^ T cells, (**d**) NK cells, (**e**) neutrophils, (**f**) total monocytes, (**g**) Ly6C^high^, Ly6C^low^, and Ly6C^-^ monocytes. Data are presented as box and whisker plots, minimum to maximum, with the center line at the median, and each data point indicates an individual mouse. CTL (black circles) n=10, HC (grey squares) n=7. Statistical significance was assessed by two-tailed Student’s t test with Welch’s correction for unequal variances or Mann-Whitney U test according to data normality. *p<0.05.

**Supplementary Figure 7.**
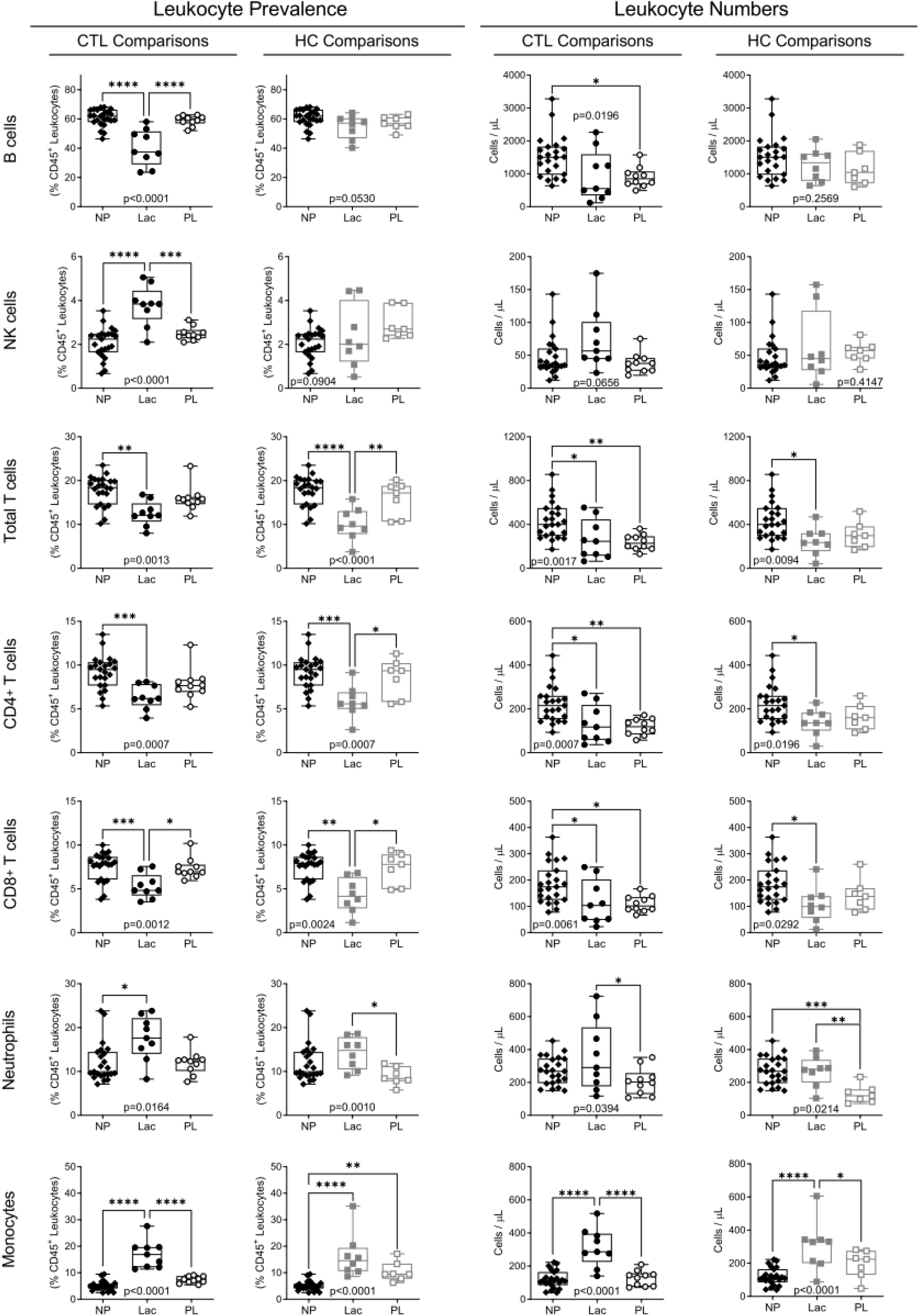
Temporal changes to maternal peripheral blood immunophenotype. Comparisons between maternal assessments of peripheral blood immunophenotype of mice in lactation (Lac) or post-lactation (PL) fed chow (CTL) or high calorie (HC) diet (from Figures 6 and 8) and age-matched never-pregnant virgin female mice fed chow diet (NP). Data are presented as box and whisker plots, minimum to maximum, with the center line at the median, and each data point indicates an individual mouse. NP (black diamonds) n=23; CTL Lac (black circles) n=9; CTL PL (open black circles) n=10; HC Lac (grey squares) n=8; HC PL (open grey squares) n=7. Statistical significance was assessed for each immune cell population by one-way ANOVA with Tukey’s multiple comparisons correction or Kruskal-Wallis test with Dunn’s multiple comparisons correction. *p<0.05, **p<0.01, ***p<0.001, ****p<0.0001.

**Supplementary Figure 8.**
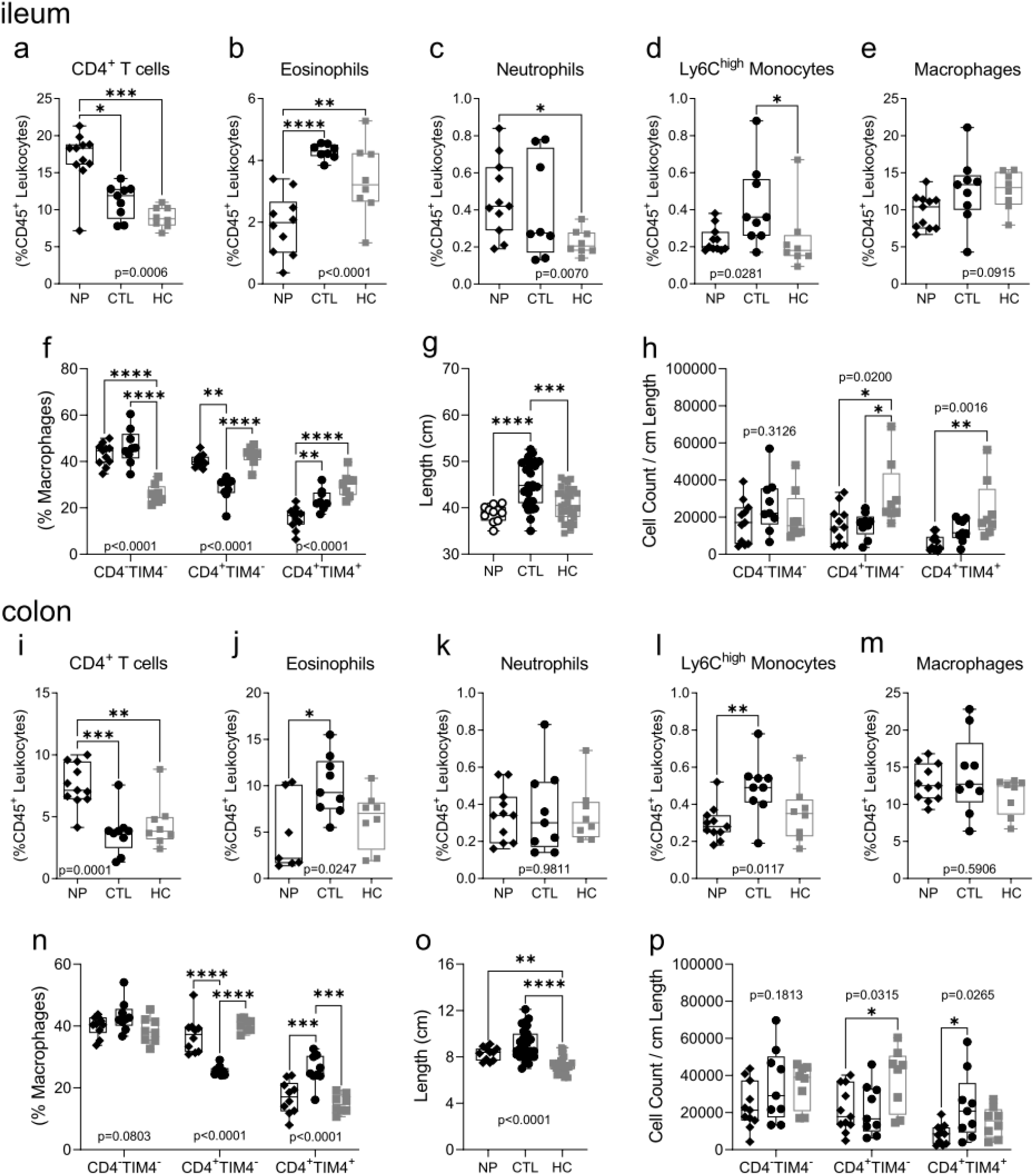
Temporal changes to maternal intestinal immune cell composition. Comparisons between maternal assessments of mice in lactation fed chow (CTL) or high calorie (HC) diet (from Figure 5 and Supplementary Figure 4) and age-matched never-pregnant virgin female mice fed chow diet (NP). Intestinal immune cell populations were assessed in ileum (**a**-**h**) and colon (**i-p**) tissues. Ileum prevalence (as a proportion of CD45^+^ leukocytes) of: (**a**) CD4^+^ T cells, (**b**) eosinophils, (**c**) neutrophils, (**d**) Ly6C^high^ monocytes, (**e**) total macrophages. (**f**) CD4^-^ TIM4^-^, CD4^+^TIM4^-^ and CD4^+^TIM4^+^ macrophages as a proportion of total macrophages. (**g**) small intestine length. (**h**) CD4^-^TIM4^-^, CD4^+^TIM4^-^ and CD4^+^TIM4^+^ macrophage cells per tissue length. Colon prevalence (as a proportion of CD45^+^ leukocytes) of: (**i**) CD4^+^ T cells, (**j**) eosinophils, (**k**) neutrophils, (**l**) Ly6C^high^ monocytes, (**m**) total macrophages. (**n**) CD4^-^TIM4^-^, CD4^+^TIM4^-^ and CD4^+^TIM4^+^ macrophages as a proportion of total macrophages, (**o**) colon length. (**p**) CD4^-^TIM4^-^, CD4^+^TIM4^-^ and CD4^+^TIM4^+^ macrophages per tissue length. Data are presented as box and whisker plots, minimum to maximum, with the center line at the median, and each data point indicates an individual mouse. NP (black diamonds) n=23, CTL (black circles) n=9, HC (grey squares) n=8. Statistical significance was assessed for each immune cell population by one-way ANOVA with Tukey’s multiple comparisons correction or Kruskal-Wallis test with Dunn’s multiple comparisons correction. *p<0.05, **p<0.01, ***p<0.001, ****p<0.0001.

## Supplementary Tables

**Supplementary Table S1.** Antibodies for surface immunophenotyping.

| Antibody | Fluorophore | Company, Catalogue # (Clone) | RRID | Dilution |
| --- | --- | --- | --- | --- |
| **Intestinal immune cell surface stain** |  |  |  |  |
| Ly6C | AF488 | BioLegend, 128022 (HK1.4) | AB_10639728 | 1:250 |
| Tim-4 | PE | BioLegend, 130005 (RMT4-54) | AB_1227807 | 1:67 |
| CD64 | PE-Dazzle 594 | BioLegend, 139320 (X54-5/7.1) | AB_2566559 | 1:200 |
| CD11b | PerCP-Cy5.5 | eBioscience, 45-0112-82 (M1/70) | AB_953558 | 1:200 |
| CCR2 | BV 421 | BioLegend, 150605 (SA203G11) | AB_2571913 | 1:100 |
| CD45 | BV 510 | BioLegend, 103138 (30-F11) | AB_2563061 | 1:200 |
| F4/80 | BV 605 | BioLegend, 123133 (BM8) | AB_2562305 | 1:100 |
| CD4 | APC | BioLegend, 100516 (RM4-5) | AB_312719 | 1:67 |
| MHC II | AF700 | eBioscience, 56-5231-82 (M5/114.15.2) | AB_494009 | 1:200 |
| CD3 | PE-Cy7 | eBioscience, 25-0031-81 (145-2C11) | AB_469571 | 1:200 |
| B220 | PE-Cy7 | eBioscience, 25-0452-81 (RA3-6B2) | AB_469626 | 1:200 |
| Ly6G | PE-Cy7 | eBioscience, 25-0112-81 (M1/70) | AB_469587 | 1:200 |
| Live-Dead NIR | APC-Cy7 | Invitrogen, L34975 | NA | 1:1000 |
| **Peripheral blood immune cell surface stain – monocytes and neutrophils** |  |  |  |  |
| CD3 | AF 700 | eBioscience, 56-0032-82, (17A2) | AB_529507 | 1:750 |
| CD11b | PE-Cy7 | eBioscience, 25-0112-82, (M1/70) | AB_469588 | 1:1200 |
| CD19 | AF 700 | eBioscience, 56-0193-82, (eBio1D3) | AB_837083 | 1:750 |
| CD45 | eFluor 450 | eBioscience, 48-0451-82, (30-F11) | AB_1518806 | 1:440 |
| Ly6G | AF 700 | eBioscience, 56-9668-82, (1A8) | AB_2802355 | 1:1500 |
| NK1.1 | AF 700 | eBioscience, 56-5941-82, (PK136) | AB_2574505 | 1:1500 |
| Ly6C | AF 488 | BioLegend, 128022, (HK1.4) | AB_10639728 | 1:1200 |
| CCR2 | PE | R&D Systems, FAB5538P, (475301) | AB_10718414 | 1:375 |
| CX3CR1 | BV650 | BioLegend, 149033, (SA011F11) | AB_2565999 | 1:1500 |
| F4/80 | APC | eBioscience, 17-4801 (BM8) | AB_2784648 | 1:1500 |
| **Peripheral blood immune cell surface stain – B cells, NK cells, T cells** |  |  |  |  |
| CD3 | BV 605 | BioLegend, 100237, (17A2) | AB_2562039 | 1:300 |
| CD4 | APC-eFluor780 | eBioscience, 47-0042-82, (RM4-5) | AB_1272183 | 1:750 |
| CD8 | APC | eBioscience, 17-0081-82, (53-6.7) | AB_469335 | 1:750 |
| CD11b | PE-Cy7 | eBioscience, 25-0112-82, (M1/70) | AB_469588 | 1:1200 |
| CD19 | AF 700 | eBioscience, 56-0193-82, (eBio1D3) | AB_837083 | 1:750 |
| CD45 | eFluor 450 | eBioscience, 48-0451-82, (30-F11) | AB_1518806 | 1:440 |
| NK1.1 | PE | eBioscience, 12-5941-82, (PK136) | AB_466050 | 1:750 |

Dilutions are calculated for a total staining volume of 50 $\mu$ L for intestinal immune cells, and 150 $\mu$ L for whole blood and bone marrow immune cells (50 $\mu$ L antibody stain with 100 $\mu$ L sample).

**Supplementary Table S2.** Summary of offspring characteristics.

|  | CTL |  | HC |  |
| --- | --- | --- | --- | --- |
|  | Female | Male | Female | Male |
| Pups per Litter |  |  |  |  |
| N (litters) | 34 |  | 39 |  |
| Mean (SD) | 3.4 ± 1.4 | 4.1 ± 1.6 | 3.8 ± 1.6 | 4.1 ± 1.5 |
| Sex Ratio (female: male) |  |  |  |  |
| Mean (SD) | 1.1 ± 1.0 |  | 1.1 ± 0.6 |  |
| Pup Weight - P7 (grams) |  |  |  |  |
| N | 34 | 33 | 38 | 39 |
| Mean (SD) | 3.83 ± 0.53 | 3.85 ± 0.43 | 3.80 ± 0.50 | 3.87 ± 0.50 |
| Pup Weight - P14 (grams) |  |  |  |  |
| N | 34 | 33 | 37 | 38 |
| Missing | 0 | 0 | 1 | 1 |
| Mean (SD) | 7.03 ± 0.75 | 6.95 ± 0.54 | 7.73 ± 0.66 | 7.78 ± 0.59 |
| Pup Weight - P21 (grams) |  |  |  |  |
| N | 34 | 33 | 38 | 39 |
| Mean (SD) | 8.85 ± 1.35 | 9.22 ± 1.55 | 10.53 ± 0.88 | 10.76 ± 0.87 |

